# Mechanism of allosteric activation in human mitochondrial ClpP protease

**DOI:** 10.1101/2024.09.27.615468

**Authors:** Monica M. Goncalves, Adwaith B. Uday, Taylor J. B. Forrester, S. Quinn W. Currie, Angelina S. Kim, Yue Feng, Yulia Jitkova, Algirdas Velyvis, Robert W. Harkness, Matthew S. Kimber, Aaron D. Schimmer, Natalie Zeytuni, Siavash Vahidi

**Author notes:** These authors contributed equally to this work (Monica M. Goncalves, Adwaith B. Uday, Taylor J. B. Forrester). **Correspondence to:** Siavash Vahidi; or Natalie Zeytuni; or Aaron D. Schimmer; or Matthew S. Kimber; or Robert W. Harkness.

## Abstract

Human ClpP protease contributes to mitochondrial protein quality control by degrading misfolded proteins. ClpP is overexpressed in cancers such as acute myeloid leukemia (AML), where its inhibition leads to the accumulation of damaged respiratory chain subunits and cell death. Conversely, hyperactivating ClpP with small-molecule activators, such as the recently-discovered ONC201, disrupts mitochondrial protein degradation and impairs respiration in cancer cells. Despite its critical role in human health, the mechanism underlying the structural and functional properties of human ClpP remain elusive. Notably, human ClpP is paradoxically activated by active-site inhibitors. All available structures of human ClpP published to date are in the inactive compact or compressed states, surprisingly even when ClpP is bound to an activator molecule such as ONC201. Here, we present the first structures of human mitochondrial ClpP in the active extended state, including a pair of structures where ClpP is bound to an active-site inhibitor. We demonstrate that amino acid substitutions in the handle region (A192E and E196R) recreate a conserved salt bridge found in bacterial ClpP, stabilizing the extended active state and significantly enhancing ClpP activity. We elucidate the ClpP activation mechanism, highlighting a hormetic effect where sub-stoichiometric inhibitor binding triggers an allosteric transition that drives ClpP into its active extended state. Our findings link the conformational dynamics of ClpP to its catalytic function and provide high-resolution structures for the rational design of potent and specific ClpP inhibitors, with implications for targeting AML and other disorders with ClpP involvement.

**Significance statement:** Human ClpP protease is essential for maintaining mitochondrial protein quality by degrading damaged proteins. In cancers like acute myeloid leukemia (AML), ClpP is overexpressed, and inhibiting it causes cancer cell death by disrupting mitochondrial function. Conversely, activating ClpP with small molecules, such as ONC201, also leads to cancer cell death by impairing mitochondrial respiration. However, the structural details of ClpP activation have been elusive. Our research presents the first structures of human ClpP in its active state, revealing a novel activation mechanism where inhibitors unexpectedly trigger activity through allosteric changes. These insights provide a foundation for designing targeted therapies for AML and other diseases where ClpP plays a crucial role.

## Introduction

ClpXP is an evolutionarily conserved proteolytic complex, consisting of the AAA+ (ATPases Associated with diverse cellular Activities) ClpX unfoldase and the ClpP protease (1). Mitochondria lack the ubiquitin-proteasome system, relying instead on proteases like ClpXP for protein quality control (2–4). ClpP is overexpressed in various cancers, including acute myeloid leukemia (AML) (5–8). Small-molecule ClpP activators like ONC201 (Dordaviprone) bind to ClpP, displace the ClpX unfoldase, and induce dysregulated mitochondrial protein degradation (9–12). ONC201 has shown preclinical efficacy in various cancers and promising results in clinical trials (13, 14). Moreover, mutations in mitochondrial ClpP have been linked to Perrault syndrome (15), an autosomal recessive disorder characterized by hearing loss and underdeveloped or absent ovaries (16). Interestingly, while some Perrault syndrome mutations lead to a loss of function, others activate ClpP (17), illustrating the complex relationship between mutations and their impact on human ClpP structure and function (18, 19).

Bacterial homologs of ClpP have been extensively studied using structural tools, revealing a stacked pair of homoheptameric rings in a barrel-like assembly (20). ClpP can adopt three distinct conformational states: extended (21–24), compact (20, 25, 26), and compressed (24, 27, 28). These are primarily distinguished by the conformation of the handle domains which in turn affects the height of the ClpP barrel (29, 30). The handle domains, consisting of a strand-turn-helix motif, mediate the interlocking of the heptameric rings of ClpP (21). The positioning of the catalytic triad relative to the handle domain directly links ClpP function to the adopted conformation of the ClpP barrel (23, 26). Only the tetradecameric extended state has the catalytic triad orientation required for activity (24). The compact and compressed states, characterized by a shortened or kinked handle helix, respectively, and a disordered handle β-strand, are inactive because the handle domain conformation misaligns the catalytic triad (31).

The functional dynamics of human ClpP remain less understood compared to its well-studied bacterial counterparts. Unlike most bacterial homologs which form stable tetradecamers, human ClpP exists as a mixture of heptamers and tetradecamers in solution (32, 33). To date, all of the available structures of human ClpP are in the inactive compact or compressed states, surprisingly even when ClpP is bound to an activator molecule such as ONC201 (Fig. 1) (10, 11, 34–39). Structures of human ClpP bound to an identifiable active-site inhibitor have not been reported yet. The absence of a structure for the active extended state of human ClpP has hampered the structure-guided design and optimization of specific active-site inhibitors similar to those developed for the proteasome (40, 41). Paradoxically, proteasome inhibitors such as bortezomib and carfilzomib initially induce significant *activation* of human ClpP before ultimately inhibiting it in mid-micromolar range (40), though the underlying mechanism remains unknown.

**Fig. 1.**
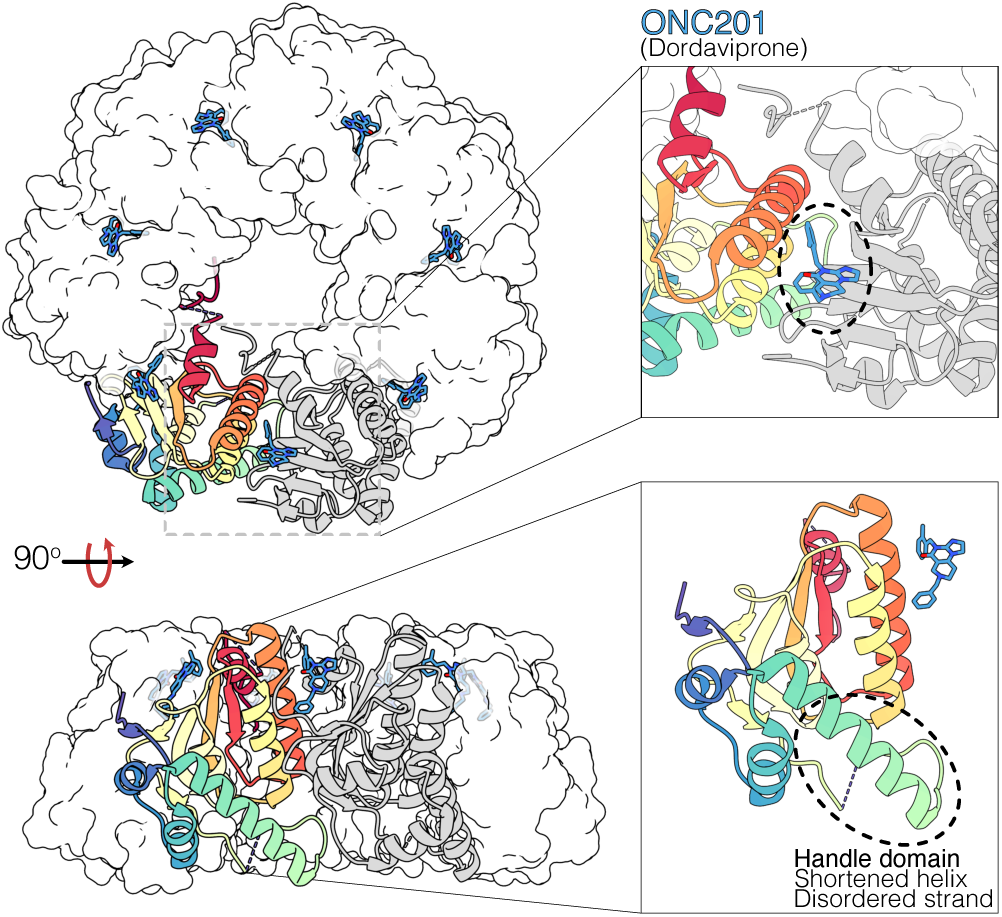
Structure of human mitochondrial ClpP bound to small-molecule activator, ONC201 (Dordaviprone) (PDB: 6DL7). ONC201 binds to the hydrophobic pockets of ClpP (inset – top) and induces dysregulated mitochondrial protein degradation. ONC201-bound ClpP is in the inactive compact state, as indicated by shortened handle domain helices and disordered handle strands (inset – bottom). The heptameric assembly in the asymmetric unit is displayed.

Here we present the first series of structures of human ClpP in the active extended state and demonstrate an allosteric mechanism by which active-site inhibitors activate human ClpP. We leverage these insights to design a pair of amino acid substitutions in the handle domain that stabilize the active extended state of ClpP, resulting in a dramatic activation of the protease. Our results establish the relationship between the conformational dynamics and catalytic activity of human mitochondrial ClpP and pave the way for the structure-guided development of potent and specific ligands against it.

## Results

### Human ClpP is activated by sub-stoichiometric binding of active-site inhibitors

Peptidyl boronates, such as bortezomib and ixazomib, are clinically-approved human proteasome inhibitors that also inhibit human ClpP, albeit with substantially lower potency (40). Here, we used these compounds to understand how active-site protease inhibitors impact the enzymatic activity and conformation of wild-type ClpP (ClpP_WT_). Similar to a previous report (40), titration with bortezomib (Fig. 2A) and ixazomib (Fig. 2B) showed a dramatic *activation* in the early phase of the dose-response curves of human ClpP_WT_ with acetyl-Trp-Leu-Ala-7-amino-4-methylcoumarin (Ac-WLA-AMC) as substrate. For example, bortezomib at 10 µM activated ClpP_WT_ approximately 20-fold, whereas ixazomib at 4 µM achieved a 30-fold activation. The degree of activation gradually diminished at higher inhibitor concentrations, as would be expected for classical competitive inhibition. These dose-response curves fit well to a hybrid model (SI Appendix) which contains terms for both activation (including a Hill cooperativity parameter *n*) as well as inhibition by the ligand. This yielded Hill coefficients of 1.9 ± 0.2 and 1.2 ± 0.25 for the activation phase of the curve (Fig. 2A and B, SI Appendix – Fig. S1). Size-exclusion chromatography (SEC) showed that apo (unbound) ClpP_WT_ elutes predominantly as a heptamer (SI Appendix – Fig. S2A), consistent with previous work (32). The addition of 10-fold molar excess of bortezomib or ixazomib results in an almost complete shift towards tetradecameric ClpP (Fig. 2C, SI Appendix – Fig. S2B). Together, these results indicate a dynamic system where ClpP shifts from an inactive heptameric state to an active tetradecameric state upon the binding of substrate-like boronates at the active site—an unexpected outcome for ligands presumed to exclusively bind to the active site of the protease.

**Fig. 2.**
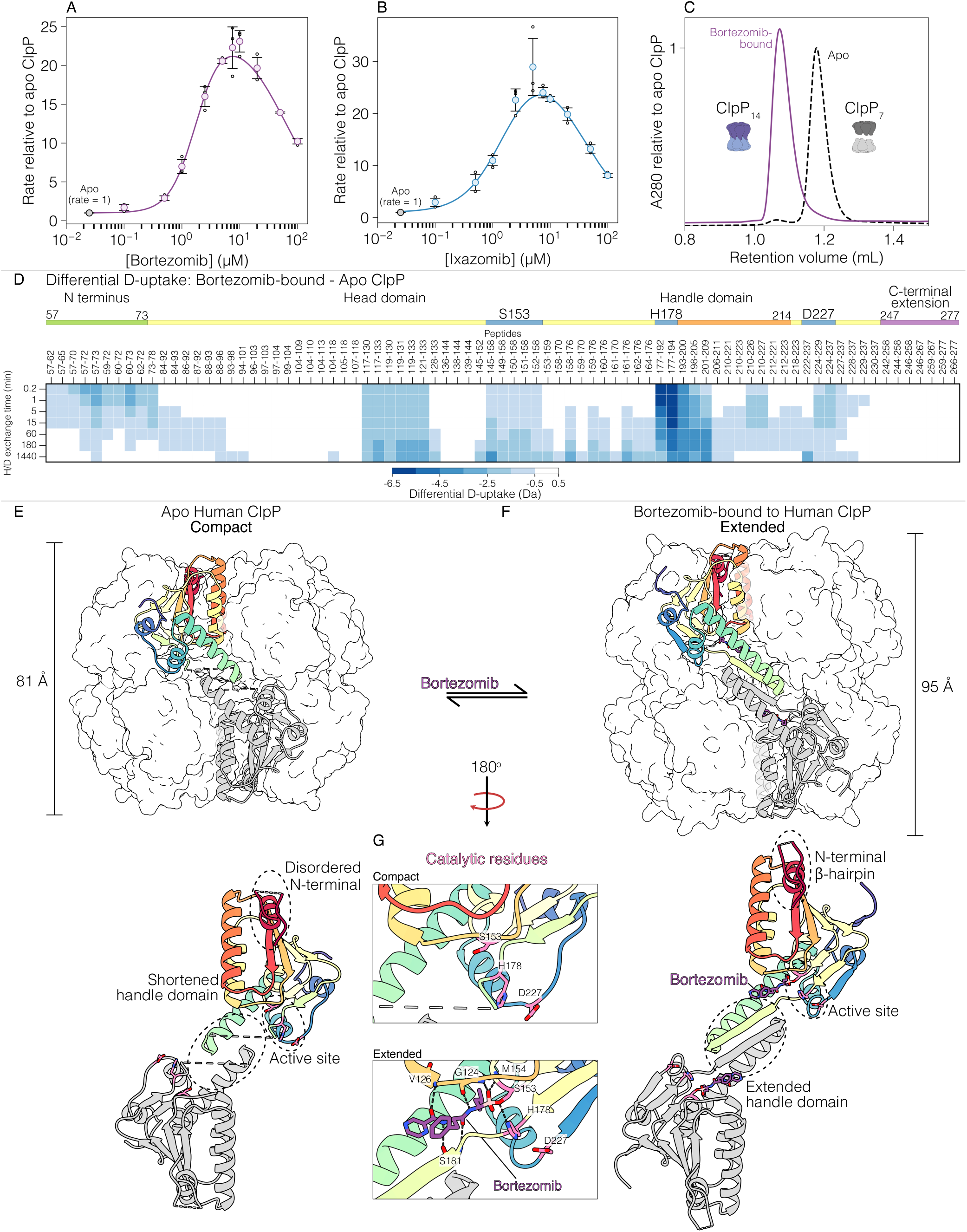
Activation of human ClpP by active-site inhibitors. Dose-response curves of human ClpP_WT_ titrated with (A) bortezomib and (B) ixazomib reveal activation at sub-stoichiometric concentrations. Initial reaction rates relative to apo ClpP_WT_ were fit to a modified Hill model (SI Appendix) which yielded Hill coefficients of 1.9 ± 0.2 and 1.2 ± 0.25, respectively; (C) Size-exclusion chromatography shows that apo ClpP_WT_ (black dashed trace) is predominantly heptameric, and the addition of 10-fold molar excess of bortezomib (purple trace) results in a shift towards the tetradecameric oligomeric state; (D) A heatmap displaying the differential deuterium uptake of bortezomib-bound ClpP_WT_ relative to apo ClpP_WT_. Bortezomib binding leads to a substantial reduction in deuterium uptake in the handle and N-terminal domains consistent with an activating structural transition. Cryo-EM structure of (E) apo and (F) bortezomib-bound ClpP_WT_ in the compact and extended states, respectively. In each case, the height of the ClpP barrel is indicated and a pair of ClpP monomers from opposing heptamers (rainbow and grey) are coloured to highlight the inter-ring interaction. The structural differences of the N-terminal domain, handle domain and catalytic residues are indicated. The dashed lines correspond to the residues which were not modelled due to lack of density; (G) Catalytic residues in apo – compact and bortezomib-bound – extended states. In the bortezomib-bound structure, the side chain of catalytic S153 is covalent bound to the boronate group of bortezomib and hydrogen bonds H178 side chain. The boronate hydroxyl group is coordinated by G124 and M154. The peptidic backbone of bortezomib hydrogen bonds with S181 of the handle β-strand, G124 and V126 of the head domain.

### Active-site inhibitor binding rigidifies the handle domain of human ClpP

We next used hydrogen deuterium exchange mass spectrometry (HDX-MS) (42) to probe the conformational dynamics of human ClpP_WT_ in response to bortezomib binding. HDX-MS is highly sensitive to molecular motions, as the HDX rate of individual backbone amides is influenced by their solvent exposure and participation in hydrogen bonding networks (43). Rigid protein segments undergo HDX slowly, while regions that undergo local conformational fluctuations become deuterated faster (44, 45). HDX-MS is especially useful as part of comparative experiments, such as when assessing the differences in conformational dynamics between the ligand-free and ligand-bound forms of a protein (46). Regions involved in ligand binding typically show a decrease in deuterium uptake, while allosteric changes in the protein can result in either an increase or decrease in uptake (47). We used these properties in a detailed structural investigation of the activation mechanism of ClpP_WT_.

Our HDX-MS workflow yielded 81 peptides with a sequence coverage of 95% and a peptide redundancy level of 4.4 (SI Appendix – Fig. S3 and Table S1). We measured the deuterium uptake of apo and bortezomib-bound ClpP_WT_ at seven time-points over a 24-hour period. We visualized our HDX-MS results as heat maps that display differential deuterium uptake for bortezomib-bound ClpP_WT_ compared to apo ClpP_WT_, as a function of D_2_O exposure time (Fig. 2D). All peptides in these two states display EX2 kinetics, characterized by fast-timescale hydrogen bond opening/closing events relative to the chemical exchange rate (*k*_ch_) (42). Bortezomib binding results in a substantial reduction in deuterium uptake across all ClpP domains, with the largest differences occurring in the active site, handle, and N-terminal regions. As expected, peptides encompassing the catalytic residues that primarily interact with bortezomib are dramatically rigidified. Most notably, the largest deuterium uptake differences are on the order of 4.5 – 7.5 Da and occur in peptides within the handle domain that encompass the catalytic histidine (residues 177 – 194). Substrates and peptidic inhibitors engage ClpP by forming hydrogen bonds with the handle β-strand (48). The HDX protection observed in these regions is consistent with these interactions. The handle helix (residues 188 – 214), which adopts multiple conformations as ClpP interconverts between the extended, compact, and compressed states, and the oligomerization sensor residues (29) (E225 and R226) are also protected (peptides 222 – 237, 224 – 229, 224 – 237) in the bortezomib-bound state (Fig. 2D).

Beyond this local effect originating from bortezomib binding and tetradecamer formation, multiple peptides in the N-terminal domain that form the axial pores (residues 57 – 73) have reduced deuterium uptake of up to 3.5 Da within the first 15 minutes of D_2_O exposure. A likely explanation for this observation is that, like its bacterial counterparts, human ClpP exhibits allosteric coupling between the N-terminal region and the active site (35, 49–51). Overall, our HDX-MS results reveal a significant structural transition in ClpP_WT_ upon bortezomib binding that affects the active site and the N-terminal axial pores.

### Structural basis for human ClpP activation by active-site inhibitors

To understand how sub-inhibitory concentrations of bortezomib can activate ClpP and impact its structure, we used single-particle electron cryo-microscopy (cryo-EM) to determine the structure of human ClpP_WT_ in the apo and bortezomib-bound forms. Two-dimensional (2D) classification of the apo ClpP_WT_ particle images yielded a mix of heptameric and tetradecameric particles (SI Appendix – Fig. S4). Following this, we performed multiple rounds of 3D classification, which gave primarily tetradecameric structures. We used 44% of the selected tetradecamer particles from the 2D classification for high-resolution refinement with enforced D7 symmetry. The final refined map had a global resolution of 2.8 Å (SI Appendix – Fig. S5). Additional classifications without symmetry enforcement did not reveal other conformations. The final map showed a tetradecameric structure consisting of two heptameric ClpP rings in the compact state. The resolution of the reconstructed map was sufficiently high to unambiguously generate molecular models (SI Appendix – Fig. S6). This structure bears strong resemblance to the ONC201-bound structure of human ClpP (C_α_ RMSD of 0.66 Å, calculated per subunit). It is characterized by the absence of density for the handle β-strand (residues 181 – 187), a handle helix truncated by two turns (residues 195 – 214) relative to the extended state, and a reduced barrel height of 81 Å (Fig. 2E). The active extended structure of ClpP is stabilized by an electrostatic network involving the oligomerization sensor residues and the conserved HQP motif which includes the catalytic H178. In our structure of apo ClpP_WT_, the oligomerization sensor residues (29) (E225 and R226) are close enough to interact. However, the loop containing Q179 swings away from the oligomerization sensor residues, which in turn misaligns the catalytic H178 and renders the compact state inactive (SI Appendix – Fig. S7). Approximately 32% of the particles resembled heptamers, but due to their highly dynamic nature, we were unable to obtain a high-resolution map of these assemblies. As with the tetradecameric compact state, disengagement of the oligomerization sensors in the heptameric form misaligns the catalytic triad into a conformation that is incapable of performing catalysis. Notably, none of the classes in our 3D classification are consistent with the geometry of apo ClpP_WT_ in an active extended state, reflecting the fact that this conformation represents only a small fraction of the total particles on the EM grid and in solution, consistent with the low activity of ClpP_WT_ in our functional assays.

Cryo-EM analysis of apo ClpP_WT_ in the presence of a two-fold molar excess of bortezomib resulted in predominantly tetradecameric 2D classes. After multiple rounds of 3D classification, 25% of the particles selected from the 2D classification step represented intact tetradecamers (SI Appendix – Fig. S8). We used these particles for the final refinement, yielding a reconstructed map with a global resolution of 2.5 Å (SI Appendix – Fig. S5B and S8). We observed no additional conformations upon further classification without symmetry enforcement. The refined map revealed a tetradecameric arrangement consisting of paired heptameric ClpP rings. In contrast to apo ClpP_WT_, bortezomib-bound ClpP was in the active extended state, evidenced by a fully formed handle helix and strong density for the handle β-strand (Fig. 2F, SI Appendix – Fig. S6D-F). Akin to bacterial ClpP homologs (20, 50, 52, 53), and consistent with our HDX-MS results showing decreased deuterium uptake (Fig. 2D), the N-terminal residues form a β-hairpin in the bortezomib-bound extended state. Clear density for bortezomib shows it in a position consistent with a covalent bond between the catalytic serine and the boronate warhead. The catalytic S153 and H178 side chains are hydrogen-bonded, while the boronate hydroxyl group is coordinated by the backbone amide nitrogen atoms of G124 and M154. The peptidic backbone of bortezomib forms hydrogen bonds on one side with S181 on the β-strand of the handle domain, and on the other, with G124 and V126 of the head domain (Fig. 2G). The isopropyl moiety of bortezomib sits in the deep non-polar S1 pocket lined by the side chains of V126, M154, and P180, but does not reach Y209 and M224 in deeper portions of this pocket (SI Appendix – Fig. SA). Human ClpP lacks an S2 pocket, therefore the central phenylalanine moiety of bortezomib is largely exposed on the protein surface (SI Appendix – Fig. S9B). Finally, the pyrazinoic acid group sits shallowly in the S3 pocket formed by G182, I198, and L201 and interacts with the backbone carbonyl of S181 (SI Appendix – Fig. S9A). Overall, our structure reveals that the individual bortezomib side chains are sub-optimal for binding to human ClpP, highlighting the potential for developing inhibitors that more effectively accommodate the geometry of the unique substrate-binding pockets of human ClpP.

We performed 3D variability analysis (3DVA) (54) to examine continuous conformational heterogeneity in both ClpP_WT_ datasets. For apo ClpP_WT_, several components revealed a slight elongation of the ClpP barrel, accompanied by the transient appearance and disappearance of density for the oligomerization sensors (residues E225 and R226) (SI Appendix – Movie S1), consistent with our HDX-MS data, which indicate these residues are dynamic in the apo state. In the bortezomib-bound dataset, four of the six 3DVA components showed transient formation and disappearance of the oligomerization sensors and handle β-strand (SI Appendix – Movie S2 and Fig. S10), with R226 being more stabilized compared to the apo state. The remaining two components displayed flexibility in the N-terminal β-hairpin (SI Appendix – Movie S3).

### Restoring the handle helix salt bridge slows human ClpP tetradecamer dissociation

Many bacterial ClpP homologs exist predominantly in the active extended tetradecameric state. Structural studies reveal a conserved pair of complementary charged residues towards the N terminus of the handle domain where the typical *i*, *i* + 4 arrangement of side chains creates a salt bridge stabilizing this segment of the handle helix. For example, in *Neisseria meningitidis* ClpP (NmClpP) these residues are E143 and R147 (SI Appendix – Fig. S11A), which form an intra-helix interaction that appears to promote handle helix stability (29, 53, 55). Interestingly, R147 also mediates interactions that contribute to the compact state, suggesting that this position may be a key determinant of the relative stability of these two states (SI Appendix – Fig. S11B). Human ClpP_WT_ contains A192 and E196 in the equivalent positions, which precludes the formation of the salt bridge that stabilizes the extended state. In the compact state of human ClpP, E196 in a given subunit interacts with the opposite ring by forming a hydrogen bond with Y206, and a salt bridge with K202 from a subunit in the opposite ring (SI Appendix – Fig. S11C) (56). Replacing E196 would destabilize the compact state due to loss of hydrogen bonds, electrostatic repulsion with K202, and steric exclusion, as in NmClpP. We reasoned that reverting these residues to the consensus sequence would promote α-helix formation and reinforce the stability of the handle domain, with the net effect of promoting human ClpP_WT_ activation by favouring the tetradecameric extended state.

SEC revealed that mutations in the handle region of ClpP (A192E / E196R; hereafter referred to as ClpP_extended_) significantly shifted the structure towards the tetradecameric form compared to ClpP_WT_ (SI Appendix – Fig. S12). We used continuous HDX-MS to compare the conformational dynamics of ClpP_extended_ to ClpP_WT_. Similar to ClpP_WT_, ClpP_extended_ exhibited EX2 kinetics across all peptides (42). The differential deuterium uptake of apo ClpP_extended_ compared to apo ClpP_WT_ shows rigidification on a peptide spanning the handle domain (e.g. residues 177 – 194), and allosteric rigidification within the N-terminal domain (SI Appendix – Fig. S13). This is consistent with SEC results showing only a partial shift towards tetradecamer. Comparing the bortezomib-bound states of ClpP_extended_ and ClpP_WT_ that already are predominantly tetradecameric, we found that stabilization of the N-terminal and handle domains was more pronounced in the mutant background, which implies some degree of additivity between the effects of the A192E / E196R mutations and bortezomib binding (Fig. 3A). Therefore, the A192E/E196R mutations induce changes that mirror those caused by bortezomib binding, albeit with a smaller magnitude.

**Fig. 3.**
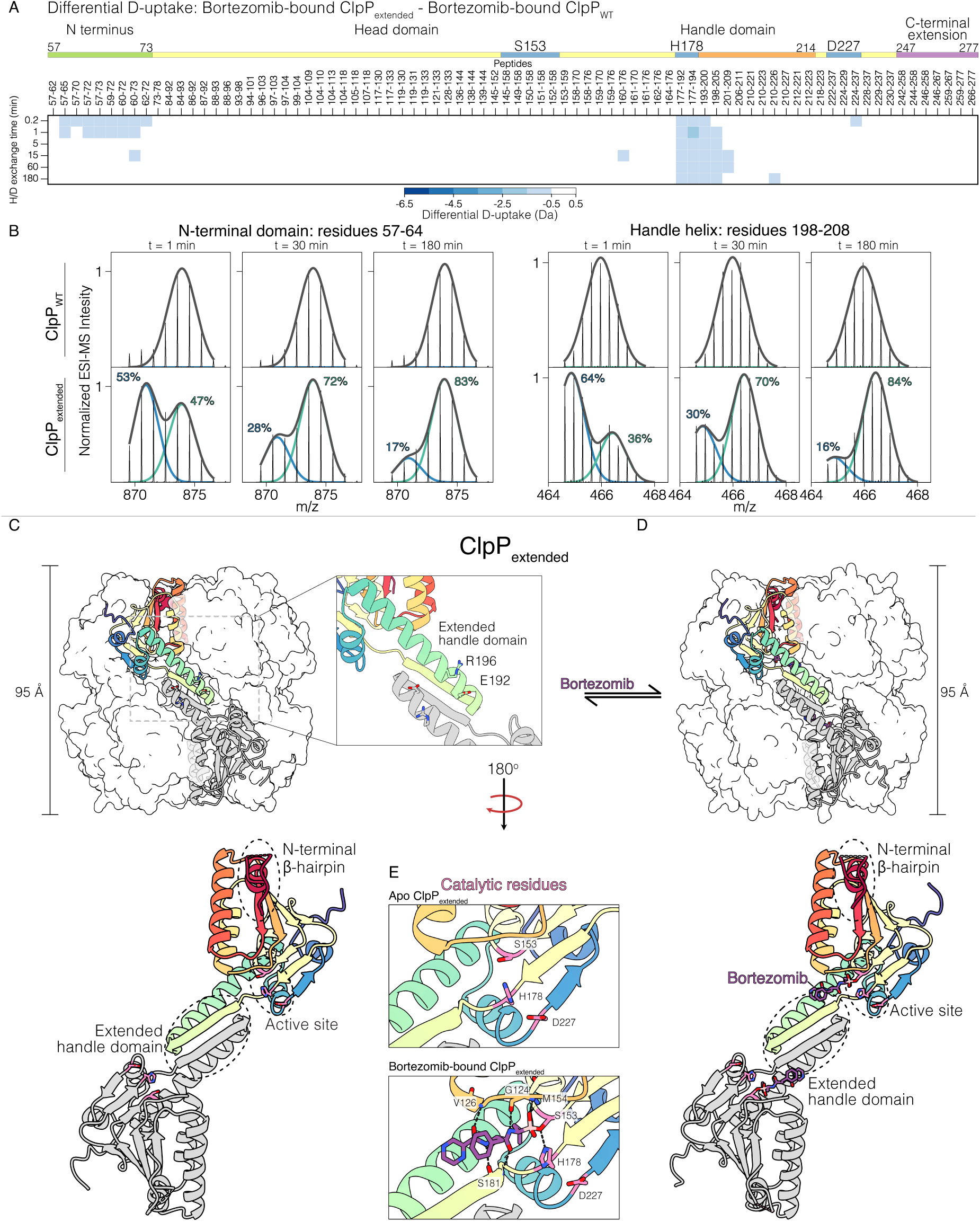
Restoring a conserved handle helix salt bridge activates human ClpP. (A) A heatmap compares the deuterium uptake of bortezomib-bound ClpP_extended_ relative to bortezomib-bound ClpP_WT_ highlighting the stabilization of the N-terminal and handle domains of ClpP_extended_. (B) Pulse HDX-MS also captures the stabilization of the tetradecameric state of ClpP_extended_. The mass spectra of N-terminal and handle domain peptides of this state are bimodal, with a protected isotopic population that declines overtime, consistent with tetradecamer dissociation. (C) The crystal structure of both apo and (D) bortezomib-bound ClpP_extended_ shows the ClpP barrel in the active and extended state, with fully formed handle domain motifs and N-terminal β-hairpins. (E) Catalytic S153 is covalent bound to bortezomib, while β-strand and head domain residues engage in hydrogen bond contacts with the inhibitor’s peptidic backbone comparable to the structures of ClpP_WT._

We used pulsed HDX-MS to measure the impact of the A192E / E196R mutations on the ClpP tetradecamer dissociation rate. We concentrated apo ClpP_WT_ and apo ClpP_extended_ to 250 µM to promote tetradecamer formation. At several intervals over a 3-hour timespan following a 100-fold dilution to start dissociation into heptamers, the ClpP samples were subjected to a 10-second D_2_O pulse. Here the mass spectra for the N-terminal and handle domain peptides of ClpP_extended_ were bimodal and were initially dominated by a protected isotopic population. The protected isotopic population decreased from 50-60% one minute after ClpP dilution to 15-20% after three hours, indicating slow dissociation of tetradecamers into heptamers (Fig. 3B). By contrast, ClpP_WT_ peptides displayed a consistent and unimodal distribution within one minute, indicating a rapid dissociation of ClpP tetradecamers to heptamers upon dilution. Taken together, our data demonstrate the functional importance of the handle region in regulating the formation of the compact and extended state, the stability of ClpP, as well as the formation of heptamer and tetradecamer complexes.

### ClpP_extended_ crystalizes as an extended tetradecamer

To better understand how the handle region of ClpP influences the conformation and activation of ClpP, we determined the structure of ClpP_extended_ at 2.8 Å by X-ray crystallography. Crystals grew in the monoclinic space group P2_1_, with a complete tetradecamer in the asymmetric unit. ClpP is in the extended state (Fig. 3C), approximately 95 Å in height, and with the handle domains fully ordered (SI Appendix – Fig. S11A). The electrostatic network stabilizing the active extended state is fully formed: the oligomerization sensor residues (29) (E225 and R226) are engaged, E225 hydrogen bonds with Q179, which in turn orients H178 optimally for catalysis (SI Appendix – Fig. S7). E192 and R196 are clearly visible in the electron density map, though their terminal functional groups generally have relatively weak density. Similar to NmClpP, the side chains of these residues approach closely enough to form a hydrogen bond in about half of the chains (SI Appendix – Fig. S14B). The active site is well defined, with clear density for the catalytic triad residues S153, H178, and D227, with all residues adopting a closely similar organization across multiple chains (SI Appendix – Fig. S14C). The active site appears suitably organized for turnover, consistent with the structure being in a catalytically active state. Notably, the ClpP_extended_ structure reported here is the first human ClpP structure in a catalytically competent conformation without any active-site inhibitor bound.

We also determined the structure of ClpP_extended_ with bortezomib soaked into the active site at 3.2 Å resolution using X-ray crystallography. Bortezomib-bound ClpP_extended_ preserves the active extended state in the crystal lattice and closely resembles the apo ClpP_extended_ structure discussed above (C_α_ RMSD of 0.32 Å, calculated per subunit). It also shows a strong similarity to the bortezomib-bound ClpP_WT_ structure we determined using cryo-EM (C_α_ RMSD of 0.69 Å, calculated per subunit). Collectively, these structures provide a structural rationale for the activation of human ClpP upon the addition of active-site inhibitors and the role of handle helix residues in the formation of the extended state.

### The conserved salt bridge in the ClpP handle helix drives dimerization and an allosteric transition leading to activation

We mapped ClpP activation kinetics by monitoring the increase in fluorescence resulting from the cleavage of Ac-WLA-AMC by ClpP_WT_ and ClpP_extended_ across a range of total enzyme and substrate concentrations (2.5 to 55 μM ClpP monomer and 2.5 to 100 μM substrate, respectively). As anticipated on the basis of our SEC and HDX-MS measurements (Fig. 2C and D), the activity profiles for ClpP_WT_ were nearly flat over the entire time course (~220 minutes) for most of our reaction conditions (SI Appendix – Fig. S15), consistent with an inability to form a tetradecameric state as a result of rapid dissociation and, in turn, an handle helix and active site conformation unable to perform catalysis. In stark contrast, the activity profiles measured for ClpP_extended_, which can readily adopt a tetradecameric form (Fig. 3C and D), exhibited a more rapid increase in fluorescence over nearly the entire set of enzyme and substrate concentrations that we employed (SI Appendix – Fig. S16). For both ClpP_WT_ and ClpP_extended_ we observed substantial lag phases leading to non-linear reaction progress curves over the first 40-minute window (Fig. 4A and B – inset). Both tetradecamer formation and the transition from an inactive to an active tetradecameric state likely contribute to the observed non-linearity in the reaction progress curves (Fig. 2, Fig. 3C). This suggested that a cooperative transition in addition to tetradecamerization is involved in the formation of a catalytically competent conformation, as indicated by our prior activity measurements in the presence of inhibitors, SEC results, and HDX-MS analyses (Fig. 2A,C and D).

**Fig. 4.**
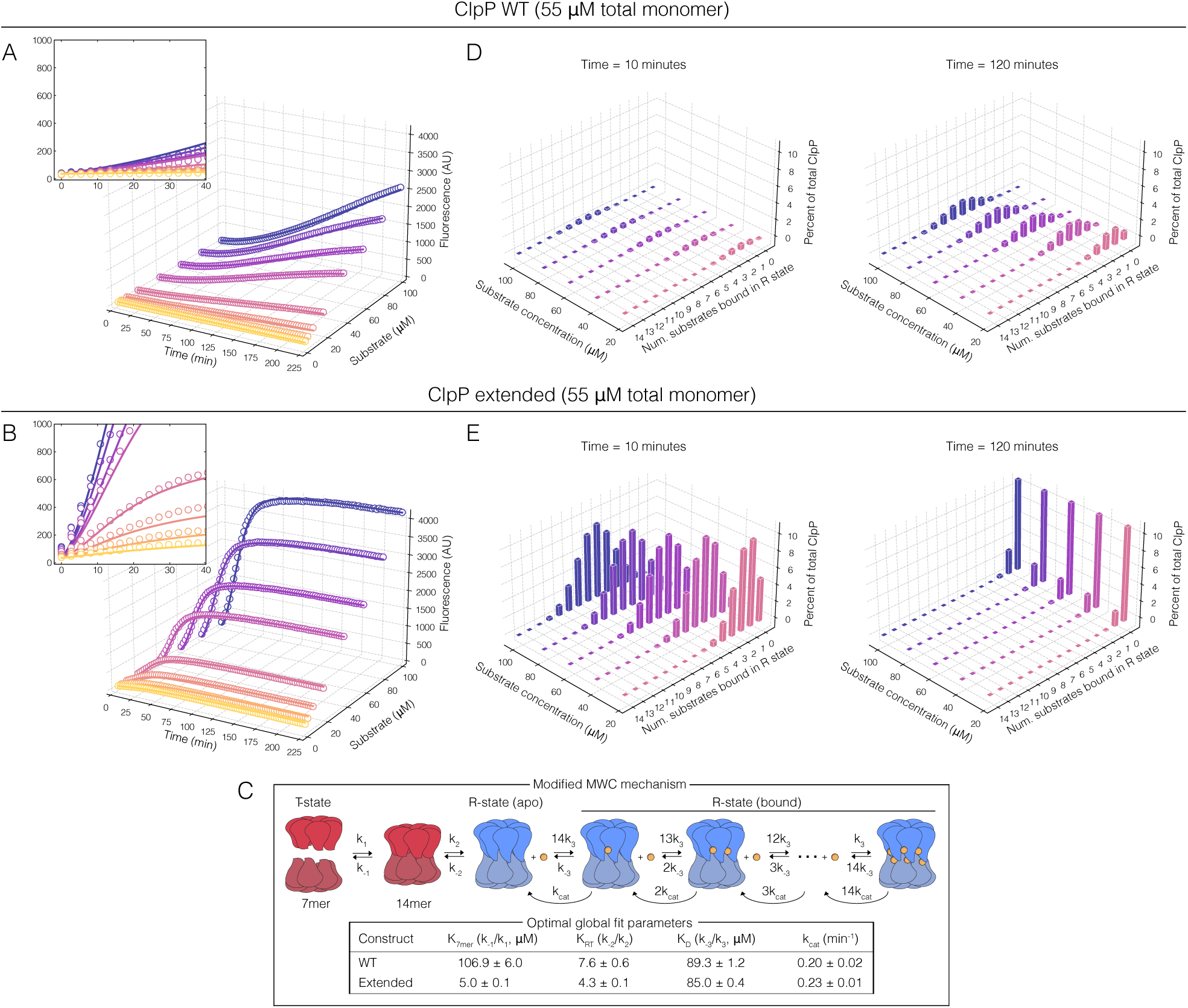
Restoration of a conserved handle helix salt bridge drives dimerization and allosteric activation of ClpP. (A and B) Peptidase activity profiles (coloured circles) for ClpP_WT_ and ClpP_extended_ (55 μM total monomer concentration) as a function of substrate concentration (2.5, 5, 10, 20, 40, 60, 80, and 100 μM) with global fits (coloured curves) of the kinetic model shown in (C). Global fits were performed with a full dataset for each enzyme leveraging total monomer concentrations of 2.5, 5, 15, 25, 35, 45, and 55 μM. To emphasize the differences between the activities of ClpP_WT_ and ClpP_extended_ only the 55 μM curves are shown here. The insets show the reaction times from 0 to 40 minutes, highlighting the sigmoidal nature of the reaction curves. (C) Modified MWC kinetic model (top) implemented in global fits of the activity profiles shown in (A, B) encompassing dimerization of ClpP heptamers (red), a classical R-T transition of apo tetradecamers (red and blue), sequential substrate binding to tetradecamers in the R state (substrates are shown as yellow circles), and substrate cleavage. The optimal parameters extracted from global fits of the data are shown in the table (bottom). Errors in the fitted parameters were estimated from Monte Carlo analyses (given as ± 2 S.D.). (D, E) Simulated percentages of (D) ClpP_WT_ and (E) ClpP_extended_ monomers within 14mers in the R conformation bound to substrates (from 0 to 14) as a function of total substrate concentration at sampled reaction times of 10 (left) and 120 (right) minutes.

To assess whether our activity data were consistent with a kinetic mechanism involving tetradecamer formation and an allosteric conformational change, we globally fit the set of time-dependent reaction profiles (14,000 data points) for each of the two enzymes to a modified MWC model (7 fitting parameters) that includes a heptamer-tetradecamer oligomeric transition and catalytic steps for substrate cleavage (Fig. 4C). We obtained good fits across the entire range of conditions in each case. These yielded equilibrium constants for ClpP tetradecamer disassembly (*K*_7mer_), a classical relaxed state (R) to tense state (T) transition of the tetradecamers (*K*_RT_), substrate dissociation from R state tetradecamers (*K*_D_), and the rate constant for peptide cleavage (*k*_cat_) (Fig. 4C). Remarkably, *K*_7mer_ for ClpP_extended_ was 21-fold smaller compared to ClpP_WT_ (5.0 ± 0.1 vs. 106.9 ± 6.0 μM). This underscores the dramatic stabilization afforded by the A192E / E196R mutations in the handle helix. In addition, *K*_RT_ was found to favour the T state by a factor of 1.8 less than for ClpP_WT_ (7.6 ± 0.6 vs. 4.3 ± 0.1 for ClpP_extended_), indicating that the identity of the handle helix residues plays an important role in the allosteric switch between active and inactive conformations, as well as in tetradecamer formation. Finally, *K*_D_ and *k*_cat_ between the two constructs were nearly identical (89.3 ± 1.2 vs. 85.0 ± 0.4 μM and 0.20 ± 0.02 vs. 0.23 ± 0.01 min^-1^ for ClpP_WT_ and ClpP_extended_, respectively), revealing that the handle helix mutations do not perturb interactions between the substrate and the active site residues.

In our model, the total pool of ClpP monomers is distributed over 17 possible oligomeric apo or ligand-bound states at any given time. To visualize the influence of handle helix on the ClpP energy landscape, we used our global fit parameters to calculate the fraction of ClpP_WT_ and ClpP_extended_ monomers within relaxed state apo and substrate-bound tetradecamers as a function of substrate concentration, assuming a total monomer concentration of 55 μM (Fig. 4D and E). At 10 minutes, ClpP_WT_ is weakly populated with substrates, while ClpP_extended_ is saturated to a much greater extent. The 120-minute time point for ClpP_WT_ revealed a larger, yet nevertheless miniscule fraction of bound tetradecamers. Unlike ClpP_WT_, the ClpP_extended_ ensemble at 120 minutes was nearly entirely shifted back toward unbound tetradecamer and heptamer as the substrate has been essentially completely converted to free product. This analysis of peptidase kinetics agrees with structural and biophysical data described above and reveals a nuanced ClpP energy landscape that governs oligomeric and allosteric structural transitions underlying its activation.

## Discussion and concluding remarks

Despite the elegant structural work conducted by several groups, human ClpP had not been captured in its active extended state. The previously determined structures of human ClpP are unsuitable for studying active site inhibitors as ClpP consistently adopts the compact state with a disorganized catalytic site and poorly resolved handle domain, surprisingly even when bound to activator molecules (10, 11, 34–39). To the best of our knowledge, the sole exception to this is a structure from 2004 (20), where the catalytic serine is attached to an unidentified ligand modeled as N-formyl methionine. The mechanism underlying the persistence of these inactive states and the activation of human ClpP upon the addition of active-site inhibitors remained unknown.

We present three structures of human ClpP in the active extended state: human ClpP_WT_ bound to bortezomib, determined using cryo-EM, and ClpP_extended_ bearing the A192E/E196R mutations that stabilize the handle region and therefore the extended state, both in the apo and bortezomib-bound states, determined using X-ray crystallography. The structures of bortezomib-bound ClpP reveal that the bortezomib side chains do not interact optimally with ClpP, which accounts for its relatively low potency. Our structural data provide a foundation for designing potent and specific boron-based peptidic inhibitors for human ClpP, akin to those developed for the proteasome. The ClpP_extended_ variant is especially useful as it significantly shifts the conformational equilibrium towards the active extended state, allowing the crystallization of human ClpP in the active extended state with an unoccupied active site. This enables substrate analogs to be soaked in, bypassing the need to co-crystallize such molecules and potentially allowing additional experiments like short soaks with reactive substrates. The consensus-restoring amino acid substitutions that stabilize the extended state are located on the solvent-exposed face of the handle region, leaving the substrate binding pockets that determine substrate specificity unaffected.

The critical role of the handle domain in regulating ClpP activity is underscored by mutations that weaken key interactions between ClpP heptamers, leading to resistance to ONC201 treatment in acute lymphoblastic leukemia cells (10). While one might expect resistance to stem from a mutation in the ONC201 binding pocket, it is instead caused by a D190A mutation in the handle region (10). In our structure of unbound ClpP_WT_, the region around D190 appears disordered, with missing density for residues 181-192. However, in our structures of the active extended form of ClpP, D190 forms a key inter-subunit salt-bridge with R226 of the oligomerization sensors and hydrogen bonds with a pair of glutamine residues at either end of the handle β-strand (Q179 and Q187) — interactions crucial for ClpP activation. Our HDX results further support this, showing that peptides that include D190 are nearly fully deuterated in the apo ClpP_WT_ state, but are significantly protected upon bortezomib binding or following the introduction of the A192E/E196R mutations that strengthen the handle domain. The D190A mutation disrupts these interactions and inactivates ClpP, even in the presence of ONC201. These findings highlight the crucial role of the handle domain in ClpP activation and emphasize its importance in mediating resistance to ClpP-targeted therapeutics.

Our study underscores the importance of carefully considering dosage and inhibitor distribution when developing therapies that target allosteric enzymes such as ClpP. Under our experimental conditions, ClpP rarely becomes saturated with substrates, even at high enzyme and substrate concentrations, indicating that there are always available sites for modulation by small molecules. This presents opportunities for targeting ClpP through either activation or classical competitive inhibition at saturating inhibitor concentrations. While potent inhibitors can effectively inhibit ClpP when bioavailable and well-distributed, low inhibitor concentrations activate the enzyme, potentially causing unintended effects. This phenomenon, known as hormesis in pharmacology (57), presents significant challenges for directing ClpP activity with substrate-mimicking compounds, and for other allosterically regulated oligomeric systems, when fine-tuning of activity may express greater biological potency than complete inhibition or activation. Modest activation effects have been observed previously, such as in *Thermus thermophilus* ClpP which exhibited a 2- to 3-fold activation in the presence of bortezomib. The 25- to 30-fold activation seen in our study of human ClpP emphasizes the dramatic differences in allosteric regulation between the homologs (58). Our model of ClpP activation by substoichiometric concentrations of substrate-mimicking inhibitors provides a framework for understanding these systems, with parallels to MAP kinase activation by BRAF inhibitors, PERK activation (59, 60), and similar effects in aspartate transcarbamoylase (61), and DegP (62). Given the broad implications of allosteric systems like ClpP in human health, elucidating the molecular mechanisms of hormesis, as demonstrated here, can inform rational strategies for the development and precise administration of therapeutics.

## Materials and Methods

Details of gene expression, protein purification, biochemical and biophysical measurements, along with data fitting and derivation of models are provided in the SI Appendix.

## Acknowledgments

M.M.G. acknowledges support from an Ontario Graduate Scholarship. A.B.U was a recipient of a Studentship Award from the Centre de Recherche en Biologie Structurale (CRBS), funded by Fonds de Recherche du Québec (Health Sector) Research Centres Grant #288558. This work was supported by the Canadian Institutes of Health Research (PJT-186334 to S.V. and A.D.S. as well as PJT-191940 to A.D.S, N.Z., and S.V.), Cancer Research Society of Canada (127241 to A.D.S, N.Z., and S.V.), Natural Sciences and Engineering Research Council (RGPIN-2020-07113 to M.S.K.), the Ontario Institute for Cancer Research with funding provided by the Ontario Ministry of Research and Innovation, the Princess Margaret Cancer Centre Foundation, and the Ministry of Long-Term Health and Planning in the Province of Ontario. A.D.S. holds the Ronald N. Buick Chair in Oncology Research. We thank Prof. M. Strauss, K. Basu, and K. Sears at the Facility for Electron Microscopy Research (FEMR) at the McGill University for their assistance with microscope operation, data collection, and computational support. FEMR is supported by the Canadian Foundation for Innovation, Quebec Government, and McGill University. MS data were recorded at the Mass Spectrometry Facility of the Advanced Analysis Centre, University of Guelph. We thank Dr. Dyanne Brewer (University of Guelph) for assistance with MS measurements.

## Author Contributions

M.M.G., A.D.S., and S.V. initiated the project; M.M.G., A.B.U., T.J.B.F., A.V., M.S.K., A.D.S., N.Z., and S.V. designed research; M.M.G., A.B.U., T.J.B.F., S.Q.W.C., A.S.K., Y.F., J.K., N.Z., and S.V. performed research; M.M.G., A.V., R.W.H., and S.V. contributed new reagents/analytic tools; M.M.G., A.B.U., T.J.B.F., R.W.H., M.S.K., N.Z., and S.V. analyzed data; M.M.G., A.B.U., T.J.B.F., R.W.H., M.S.K., A.D.S., N.Z., and S.V. wrote the paper with assistance from other others; M.S.K. supervised the X-ray crystallography studies, N.Z. supervised the electron cryomicroscopy studies; and S.V. supervised the mass spectrometry and biochemical studies.

## Data availability

Electron microscopy density maps have been deposited in the Electron Microscopy Databank (EMD-46970 and EMD-46971). Atomic models have been deposited in the Protein Databank (PDB ID 9DKV, 9DKW, 9DQK, and 9DQL). Mass spectrometry data are available from the MassIVE database as entry MSV000095957.

