## Supplementary material for "Mechanism of allosteric activation in human mitochondrial ClpP protease": SI Appendix

**Correspondence to:**

**Classifications:**

Biological Sciences - Biochemistry

### Materials and methods

#### Plasmids and constructs

Codon-optimized genes encoding wild type human ClpP (Uniprot entry: Q16740) or the A192E and E196R variant without the mitochondrial targeting sequence (residues 1-56) were synthesized and cloned at the NcoI and BamHI restriction sites of pET24a+ vectors (Genscript and Twist). A cleavable N-terminal $His_6$-SUMO tag was incorporated into both plasmids.

#### Protein expression and purification

Plasmids were transformed into competent T7 Express *Escherichia coli* cells by heat-shock and were grown in lysogeny broth (LB) medium supplemented with 30 µg/mL kanamycin at 37 °C. Protein expression was induced at an $OD_{600}$ of 0.6 with 1 mM isopropyl β-d-1-thiogalactopyranoside (IPTG) and proceeded for 18 hours at 16 °C. Cells were harvested by centrifugation at 4,000 ×*g* for 15 min at 4 °C and resuspended in lysis buffer (50 mM Tris-HCl – pH 7.0, 300 mM KCl, 20 mM imidazole). The cells were lysed using an Emulsiflex-C3 high-pressure homogenizer (Avestin Inc.) and the homogenate was clarified by centrifugation at 20,000 ×*g* for 50 min at 4 °C. The supernatant was purified using a $Ni^{2+}$-charged chelating FastFlow™ Sepharose column, and the protein was eluted using the elution buffer (50 mM Tris-HCl – pH 7.0, 300 mM KCl, 500 mM imidazole). The eluted protein sample was incubated with Ulp1 protease for His-SUMO tag cleavage and was dialyzed against lysis buffer for 18 hours. The cleaved tag and remaining impurities were removed by passing the protein solution over the $Ni^{2+}$-charged column for a second time. Subsequently, the flow-through fraction was concentrated using 50 kDa MWCO Amicon Ultra-15 centrifugal filter units (Millipore) and then subjected to size exclusion chromatography (SEC) on Superdex 200 Increase 10/300 column (GL) (Millipore) in 50 mM Tris – pH 7.8, 100 mM KCl, 10 mM $Mg_2Cl$, 1 mM DTT. The concentration of the isolated ClpP sample was determined spectrophotometrically using extinction coefficients of 17,420 $M^{-1}cm^{-1}$ computed using the Expasy ProtParam web-based tool (https://web.expasy.org/protparam/).

#### SEC-MALS and analytical SEC

The oligomeric state of $ClpP_{WT}$ and $ClpP_{extended}$ was assessed with an OMNISEC multi-detector SEC system (Malvern Panalytical) at 20 °C fitted with OMNISEC RESOLVE and OMNISEC REVEAL modules. 100 $\mu$L protein samples at 1.8 mg $mL^{-1}$ were loaded onto a pair of P3000 Protein SEC columns connected in series (each at 300×8 mm, Malvern Panalytical) equilibrated in 50 mM Tris-HCl – pH 7.8, 100 mM KCl, 10 mM $MgCl_2$, 1 mM DTT at a flow rate of 1 mL $min^{-1}$. Using BSA as a standard, the molecular weight was calculated using light scattering detectors at 90° (right-angle light scattering) and 7° (low-angle light scattering).

The oligomeric transitions of ClpP were probed on an Agilent 1100 Series HPLC system (Agilent Technologies) at ambient temperature (~23 °C). 10 $\mu$L protein samples at 1.8 mg/mL were loaded onto a ACQUITY Premier Protein SEC column, (250 Å pore size × 4.6 mm × 150 mm, Waters) in 50 mM Tris – pH 7.8, 100 mM KCl, 10 mM $Mg_2Cl$, 1 mM DTT at a flowrate of 0.3 mL $min^{-1}$.

**Degradation Assays**

The peptidase activity of ClpP was measured at 25 °C with acetyl-Trp-Leu-Ala bearing a C-terminal fluorogenic 7-amino-4-methylcoumarin group (abbreviated Ac–WLA–AMC) as substrate (Biosynth). The reaction progress was measured with a FLUOstar Omega 96-well microplate reader (BMG Labtech) at $\lambda_{ex}$: 355 nm, $\lambda_{em}$: 450 nm in 50 mM Tris-HCl – pH 7.8, 100 mM KCl, 10 mM $MgCl_2$, 1 mM DTT and 4% dimethylsulfoxide (DMSO) (v/v). For inhibitor dose-response assays, ClpP was maintained at 2.5 $\mu$M (monomer basis) and substrate at 100 $\mu$M, while bortezomib or ixazomib (Selleck Chemicals) were added at the desired concentrations from DMSO-solubilized stock solutions. To allow for ClpP's slow dissociation to equilibrate, the mixtures were incubated for 2 hours before adding the substrate from a high concentration DMSO stock to start the reactions. We demonstrated that the enzymatic activity of our ClpP protein samples is entirely dependent on the catalytic S153 of human mitochondrial ClpP and not any other potential contaminant present in the protein sample. Mutating this residue to alanine abolished the degradation of Ac-WLA-AMC (SI Appendix – Fig. S17). All degradation assays were carried out in technical triplicates, and activity was normalized relative to the apo uninhibited control.

The dose-response relationship for an allosteric activator can be approximated by the Hill model. This curve starts from a minimum response, here normalized to 1, increases with each dose increment, and eventually reaches $V_{max}$:

$$V([x]) = V_{max} - \frac{V_{max} - 1}{1 + \left(\frac{[x]}{EC_{50}}\right)^n}$$

Here, $EC_{50}$ is the effective concentration (dose) of the allosteric effector generating 50% of $V_{max}$, $n$ is the Hill coefficient which quantifies the sigmoidality of the curve, and $[x]$ represents the concentrations of the allosteric ligand. A similar function can also describe inhibitory responses, where the response begins at maximum, here normalized to 1, decreases with increasing inhibitor dose $[x]$, and reaches a minimum:

$$V([x]) = \frac{1}{1 + \left(\frac{[x]}{IC_{50}}\right)}$$

Some allosteric effectors can produce a stimulatory effect at low concentrations and an inhibitory effect at higher concentrations. This interaction results in a bell-shaped dose-response which can be modeled by multiplying the equations above. Here, the first and second terms describe the ascending and descending parts of the dose-response curve where $EC_{50} < IC_{50}$:

$$V([x]) = \left[V_{max} - \frac{V_{max} - 1}{1 + \left(\frac{[x]}{EC_{50}}\right)^n}\right] \cdot \left[\frac{1}{1 + \left(\frac{[x]}{IC_{50}}\right)}\right]$$

The data for both the functional characterization and dose-response curves were fit and visualized using in-house scripts written in Python v3.8.

**HDX-MS**

Continuous labelling, bottom-up hydrogen deuterium exchange mass spectrometry was performed on four states: (i) apo WT ClpP; (ii) apo $ClpP_{extended}$, (iii) bortezomib-bound WT ClpP; and (iv) bortezomib-bound $ClpP_{extended}$. Equilibration and exchange solutions contained 50 mM Tris-HCl – pH 7.8, 100 mM KCl, 10 mM $MgCl_2$, 1 mM DTT, and 4% DMSO (v/v). $D_2O$-based solutions were adjusted to pD 7.4 using the standard electrode correction procedure (1, 2). During the equilibration step, protein stocks were diluted to 8 $\mu$M with an $H_2O$-based buffer and for the bortezomib-bound states 2 mM bortezomib was present. HDX was initiated at room temperature (22 °C) by a 20-fold dilution into a $D_2O$-based buffer that contained additives identical to those of the equilibrated mixture. This was done such that ClpP did not experience any change in bortezomib concentration upon $D_2O$ dilution. Labelling times ranged from 0.16 to 1440 minutes. Reactions were quenched by acidification to $pH_{read}$ 2.5 by mixing sample aliquots 1:1 (v/v) with 250 mM $NaH_2PO_4$ – pH 2, 3 M guanidium chloride, and 3 mM n-dodecylphosphocholine (3) and flash frozen in liquid $N_2$. Samples were stored at -80°C prior to analysis.

Reverse-phase separation was performed using a M-Class nanoAcquity UPLC system equipped with HDX technology (Waters). 10 pmol of sample was digested at 15 °C using a nepenthesin-2 column (AffiPro, AP-PC-004, 1 mm × 20 mm). The resulting peptides were trapped for 3 min on a BEH C18 (1.7 µm, 2.1 mm × 5 mm, Waters) column at a flowrate of 100 µL $min^{-1}$. Peptides were separated on an HSS T3 column (1.8 µm, 1.0 × 50 mm, Waters) using a linear, 8-minute acetonitrile:$H_2O$ gradient acidified with 0.1% formic acid (acetonitrile ramped from 5 – 35%) at 0 °C and at a flowrate of 100 µL $min^{-1}$. Between each injection, the syringe and sample loop were cleaned using 1.5 M guanidine hydrochloride, 4% (v/v) acetonitrile, 0.8% (v/v) formic acid, and 1.5 mM n-dodecylphosphocholine to minimize column carry-over.

The UPLC outflow was directed to a quadrupole ion mobility time-of-flight (Q-TOF) Synapt G2-Si mass spectrometer (Waters) fitted with a standard electrospray source operated in positive-ion mode with a capillary voltage of +3 kV. Dynamic calibration of the instrument was achieved by infusing LeuEnk solution (1+, m/z 556.2771 *Th*) from the LockSpray capillary at a flow rate of 10 µL $min^{-1}$ sampled every 20 seconds. Drift time-aligned $MS^E$ data-independent acquisition was employed for peptide mapping, as described previously (4). Data were acquired over the 50 – 2000 *m/z* range with a scan time of 0.4 sec. Fragmentation was induced by a linear increase of the transfer collision energy in alternating scans over 20 – 40 V. Ion mobility separation was controlled manually as described previously (4). The quadrupole was manually set to dwell at 300 *m/z* to exclude smaller ions from entering the mobility cell. The TOF mass analyzer was operated in resolution mode.

Undeuterated and deuterated samples were analyzed as technical replicates. $MS^E$ data were analyzed using Waters ProteinLynx Global Server (v3.0.3) to identify peptide sequences. HDX-MS data analysis was carried out using DynamX v3.0 (Waters). Peptide filtering parameters were taken from Sørensen et al (5). The amino acid substitutions did not affect the chemical HDX rate, $k_{ch}$, significantly. Monitoring the deuterium uptake of peptides that engage bortezomib was facilitated by the rapid dissociation of the inhibitor from ClpP upon acidification.(6) The spectra corresponding to the peptides that satisfied stringent filtering criteria were manually inspected, and those with high quality spectra and high signal-to-noise ratios were retained for further analysis. A hybrid significance test model was used on selected peptides with deuterium uptake differences greater than ± 0.5 Da and that also passed Welch's *t*-test with $\alpha < 0.01$ (7). Heatmaps were generated for the visualization of HDX-MS results via HDgraphiX (8).

**Commented [SV1]:** Nguyen, D., Mayne, L., Phillips M.C., & Englander, S.W.: Reference parameters for protein hydrogen exchange rates. J Am Soc Mass Spectrom. (2018)

### Cryo-electron microscopy sample preparation and data collection

Purified $ClpP_{WT}$ was concentrated to 15 mg/ml (625 $\mu$M of monomeric ClpP). For the bortezomib-bound state, the inhibitor was added at a final concentration of 1.25 mM. To prepare this mixture, 2.5 $\mu$L of 25 mM bortezomib stock (dissolved in 100% DMSO) was added to 95 $\mu$L of $ClpP_{WT}$, and the mixture was incubated at 4 °C for 1 hour to allow the inhibitor to bind. An additional 2.5 $\mu$L of bortezomib stock was subsequently added to the mixture to reach the desired final bortezomib concentration. The mixture was washed to remove the excess DMSO, ensuring it would not interfere with specimen preparation and data collection. Protein sample was applied to a holey carbon grid (C-flat CF-1.2/1.3-3Cu-T), which had been glow-discharged in air at 10 mA for 15 seconds. Sample vitrification was performed using a Vitrobot Mark IV (Thermo Fisher Scientific) at 4°C and 100% humidity. Each protein sample was incubated on the grid for 3 sec and the grid was blotted for 5 sec with a blot force of 1 before plunging in liquid ethane. Datasets were collected using the SerialEM software (9) on a Titan Krios microscope (Thermo Fisher Scientific). Movies were recorded on a Gatan K3 direct electron detector equipped with a Quantum LS imaging filter. The total electron dose per movie was 60 $e^-/Å^2$, equally spread over 30 frames. Data were collected at a magnification of ×130,000, yielding images with a calibrated pixel size of 0.675 Å. The nominal defocus range used during data collection was between -1.00 $\mu$m and -2.5 $\mu$m. A summary of the data collection parameters for all datasets can be found in Supplemental Table 1.

#### Cryo-electron microscopy data processing

The cryo-EM data was initially processed in RELION-4 (10). The movies were corrected for beam-induced motion using RELION's implementation of MotionCorr2, and the CTF parameters were estimated using CTFFIND4 (11). Particles were picked from a subset of micrographs using RELION's LoG-based auto-picking with a minimum diameter of 70 Å and maximum diameter of 150 Å. These particles were subjected to two rounds of 2D classification, and the selected particles which resembled the ClpP tetradecamer or heptamer were used to train the TOPAZ neural network (12). The trained TOPAZ network was used to pick particles from the entire dataset. These picked particles were subjected to two rounds of 2D classification for particle curation. The particles from the best classes were then used for an initial model reconstruction, followed by one or two rounds of 3D classification with 5 classes each. The classes displaying the characteristic structural features of the ClpP tetradecamer were selected and exported to cryoSPARC v4 for further processing (13). These particles were subjected to reference-based motion correction in cryoSPARC to correct for beam-induced motion at the per-particle level. These motion-corrected particles were subsequently used to obtain the final high-resolution map with D7 symmetry enforced, combined with global and local CTF refinement (14). Local resolution was estimated for all the maps using cryoSPARC v4. ChimeraX (15) was used for visualizing the maps and figure preparation. Data processing details for the datasets are summarized in Figures S3 and S4. We also used cryoSPARC's 3D variability analysis (3DVA) (16) module to probe conformational heterogeneity in our datasets. The particles corresponding to the final refined maps were symmetry expanded to D7 symmetry, and the resulting particles were used as input for the 3DVA with six variability components and a low-pass filter resolution of 3.5 Å.

#### Cryo-EM model building and refinement

To build the molecular model, we used the crystal structure of human ClpP (PDB ID: 8I7X)(17) as an initial template for both the datasets. The template model was initially docked onto the maps using ChimeraX (15). For the bortezomib-bound dataset, the restraints for the serine hydroxyl-boron bond were generated using JLigand (18). Model building was performed by iterative cycles of manual building in Coot (19) and real space refinement in PHENIX (20), which substantially improved both the model's geometry and its fit to the map. Model validation was also performed on PHENIX (20), with a summary of the model statistics provided in Supplemental Table 1.

#### Crystallization and structure determination

$ClpP_{extended}$ was desalted into minimal buffer containing 20 mM Tris-HCl pH 8.0, 150 mM NaCl and concentrated to a final concentration of 15 mg/mL using a 10 kDa molecular weight cut-off Amicon Ultra Centrifugal Filter (Millipore-Sigma). Crystallization experiments were conducted in a sitting-drop format at room temperature, with protein mixed with (in a 2:1 or 1:1 ratio) and equilibrated against ~80 µL of well solution containing 200 mM potassium acetate and 35% (v/v) pentaerythritol propoxylate (5/4 PO/OH). Large, plate-like crystals generally grew within seven days. For the bortezomib-bound structure, crystals were soaked by mixing a drop of a 50 mM bortezomib stock (in 100% DMSO) into wells with mature crystals, followed by incubation at room temperature for 1 – 3 hrs. Crystals were cryoprotected by pulling the crystal through a layer of paratone-N oil laid over the drop to minimize dehydration of the crystals. Crystals were then frozen in liquid nitrogen for data collection at 100 K.

Data was collected at the Canadian Light Source Beam Line ID1. Data was processed and scaled using the XDS package (21). Both structures were determined using molecular replacement in Phaser in Phenix (22), using an edited (low confidence regions removed) AlphaFold model of a human ClpP protomer (AF-Q16740-F1) (23) comprised of residues 57 – 249. Rebuilding was performed in Coot (24), with refinement in Phenix.refine (25). Data collection and structure refinement statistics are shown in Table 1. All structure figures were prepared using ChimeraX or Pymol 2.5.4 (Schrödinger LLC).

#### Global fitting of ClpP peptidase activity profiles

Peptidase activity profiles were modelled with a modified Monod–Wyman–Changeux (MWC) mechanism (26). In the absence of substrate, it was assumed that ClpP interconverts between tense (T) inactive heptamers and tetradecamers ($T_7$ and $T_{14}$, respectively) that are substrate-binding incompetent according to

$$T_7 \underset{k_{-1}}{\overset{k_1}{\rightleftarrows}} T_{14} \qquad (1)$$

where $k_1$ and $k_{-1}$ are the forward and reverse rate constants governing the exchange between the two oligomeric states. It was further assumed that $T_{14}$ undergoes an intramolecular transition to a relaxed (R) state ($R_{14}$) that can bind and process substrates

$$T_{14} \underset{k_{-2}}{\overset{k_2}{\rightleftarrows}} R_{14} \qquad (2)$$

with forward and reverse rate constants $k_2$ and $k_{-2}$. Substrate binding to $R_{14}$ occurred sequentially, up to the maximum number of 14, through the equilibria

$$R_{14} + S \underset{k_{-3}}{\overset{14k_3}{\rightleftarrows}} R_{14}S_1$$

$$R_{14}S_1 + S \underset{2k_{-3}}{\overset{13k_3}{\rightleftarrows}} R_{14}S_2$$

$$\vdots$$

$$R_{14}S_{i-1} + S \underset{ik_{-3}}{\overset{(14-i+1)k_3}{\rightleftarrows}} R_{14}S_i \quad (3)$$

$$\vdots$$

$$R_{14}S_{13} + S \underset{14k_{-3}}{\overset{k_3}{\rightleftarrows}} R_{14}S_{14}$$

where $k_3$ and $k_{-3}$ are the microscopic substrate peptide association and dissociation rate constants respectively. The multiplicative factors for the rate constants take into account the number of ways to bind and dissociate a substrate in a given step. The rate constants for the forward and reverse processes respectively were taken to be identical in each binding equilibrium, i.e. substrate binding is non-cooperative. While it is straightforward to include a cooperativity parameter allowing for distinct substrate binding constants, this was not warranted by our data as excellent fits were obtained in the absence of cooperative binding. Finally, proteolysis of substrate peptides by each of the bound R states ($R_{14}S_i,\ i \in \{1 \ldots 14\}$) with concomitant product release occurred via irreversible steps according to

$$R_{14}S_1 \overset{k_{cat}}{\rightarrow} R_{14} + P$$

$$R_{14}S_2 \overset{2k_{cat}}{\rightarrow} R_{14}S_1 + P$$

$$\vdots$$

$$R_{14}S_i \overset{ik_{cat}}{\rightarrow} R_{14}S_{i-1} + P \qquad (4)$$

$$\vdots$$

$$R_{14}S_{14} \overset{14k_{cat}}{\rightarrow} R_{14}S_{13} + P$$

where $k_{cat}$ is the catalytic rate constant for conversion of bound substrate into products $P$ and their release from the enzyme. Like the above, the multiplicative factors for $k_{cat}$ account for the number of available substrates in a given complex, one of which is turned over in each step. The two product peptides have been subsumed into the single product variable; this simplification has no effect on the outcome of the model as both species are formed and released according to $k_{cat}$.

Equations 1-4 can be recast in the form of rate equations controlling the fluxes between each of the ClpP states

$$\frac{d}{dt}[T_7] = -2k_1[T_7]^2 + 2k_{-1}[T_{14}]$$

$$\frac{d}{dt}[T_{14}] = k_1[T_7]^2 - k_{-1}[T_{14}] - k_2[T_{14}] + k_{-2}[R_{14}]$$

$$\frac{d}{dt}[R_{14}] = k_2[T_{14}] - k_{-2}[R_{14}] - 14k_3[R_{14}][S] + k_{-3}[R_{14}S_1] + k_{cat}[R_{14}S_1] \qquad (5)$$

$$\frac{d}{dt}[R_{14}S_1] = 14k_3[R_{14}][S] - k_{-3}[R_{14}S_1] - 13k_3[R_{14}S_1][S] + 2k_{-3}[R_{14}S_2] - k_{cat}[R_{14}S_1] + 2k_{cat}[R_{14}S_2]$$

$$\vdots$$

$$\frac{d}{dt}[R_{14}S_i] = (14 - i + 1)k_3[R_{14}S_{i-1}][S] - ik_{-3}[R_{14}S_i] - (14 - i)k_3[R_{14}S_i][S] + (i + 1)k_{-3}[R_{14}S_{i+1}] - ik_{cat}[R_{14}S_i] + (i + 1)k_{cat}[R_{14}S_{i+1}]; 2 \le i \le 13$$

$$\vdots$$

$$\frac{d}{dt}[R_{14}S_{14}] = k_3[R_{14}S_{13}] - 14k_{-3}[R_{14}S_{14}] - 14k_{cat}[R_{14}S_{14}].$$

The rate equations for the substrate and product are thus given by

$$\frac{d}{dt}[S] = -k_3[S](14[R_{14}] + \sum_{i=1}^{13}(14 - i)[R_{14}S_i]) + k_{-3}\sum_{i=1}^{14} i[R_{14}S_i] \qquad (6)$$

$$\frac{d}{dt}[P] = k_{cat}\sum_{i=1}^{14} i[R_{14}S_i] \qquad (7)$$

The mass conservation equations associated with Equations 5-7, which were used to constrain the numerical procedure applied during the fitting routine, are

$$M_{T,j} = 7[T_7] + 14[T_{14}] + 14[R_{14}] + 14\sum_{i=1}^{14}[R_{14}S_i] \qquad (8)$$

$$S_{T,k} = [S] + \sum_{i=1}^{14} i[R_{14}S_i] \qquad (9)$$

where $M_{T,j}$ ($i \in \{2.5, 5, 15, 25, 35, 45, 55\}$ μM) and $S_{T,k}$ ($k \in \{2.5, 5, 10, 20, 40, 60, 80, 100\}$ μM) are the total ClpP monomer and substrate concentrations used in each reaction respectively.

The system of equations 5-9 was implemented in a numerical integration algorithm to obtain the time-dependent concentrations of each species in the model. The peptidase activity profiles were subsequently calculated assuming that the fluorescence signal is linearly proportional to the product concentration in each case

$$F_{j,k}^{observed}(t) = F_{j,k}^{initial} + m_{j,k}[P]_{j,k}(t) \qquad (10)$$

where $F_{j,k}^{observed}(t)$, $F_{j,k}^{initial}$, $m_{j,k}$, and $[P]_{j,k}(t)$ are the observed fluorescence values, the initial fluorescence values (i.e. at $t$ = 0), the scaling parameters that convert product concentrations to fluorescence, and the numerically solved product concentrations for the set of reactions. Each reaction was allowed a unique initial value and scaling factor for product concentration to adjust for experimental error. The $F_{j,k}^{initial}$ values were estimated from the initial experimental data points for $ClpP_{WT}$ and $ClpP_{extended}$, while the $m_{j,k}$ were approximated using the maximum fluorescence values corresponding to the completion of the reactions for $ClpP_{extended}$.

The data fitting procedure was performed using an in-house Python program leveraging the lmfit package (https://lmfit.github.io/lmfit-py/). After initializing the fitting routine with estimates of reasonable values, (obtained from initial data simulations) for the set of rate constants ($k_l,\ l \in \{1, -1, 2, -2, 3, -3, cat\}$), the modeling procedure consisted of three main steps. First, the time-dependent concentrations of each ClpP state, free substrate, and product were calculated using the system of equations 5-9, the set of rate constant estimates, and the set of $M_{T,j}$ and $S_{T,k}$ in a numerical integration algorithm as noted above. Second, the fluorescence profiles $F_{j,k}^{observed}(t)$ were simulated using the corresponding product concentrations and equation 10. Third, the fit quality was calculated according to the residual sum-of-squared differences (RSS) between the experimental and computed data points (a total of 14,000 experimental points were used in the fits for each of $ClpP_{WT}$ and $ClpP_{extended}$). The fit parameters were then adjusted, and the process iterated until optimal fit qualities were obtained. For $ClpP_{WT}$, the rate constants were found to be strongly linearly correlated, particularly those for the forward and reverse reactions of a given step (e.g. binding). This was presumably due to its relative inactivity in most of our reaction conditions, from which it is difficult to extract accurate estimates of the full set of model parameters. The ratios of the rate constants for each step (in other words, equilibrium constants) were, however, much better defined. The correlations between the rate constants were less prominent for $ClpP_{extended}$, likely as a result of its increased activity as compared to $ClpP_{WT}$ which enabled better estimates of these parameters. To compare the two ClpP constructs, we have therefore elected to provide equilibrium constants in Figure 4 as the best-defined parameters extracted from fits of their activity profiles, which nevertheless illustrate the key differences between them. The errors given for the parameters in the table in Figure 4 were estimated from Monte Carlo simulations (± 2 S.D.).

### SI Appendix Figures

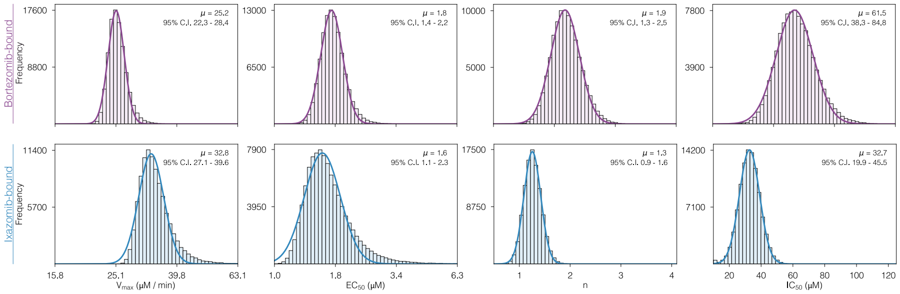


**Fig. S1.** Histograms representing 100,000 Monte Carlo simulations to determine the fitted parameters and confidence intervals for $V_{max}$, $EC_{50}$, n, and $IC_{50}$ values from the dose-response curves of human $ClpP_{WT}$ titrated with bortezomib (purple) and ixazomib (blue). The x-axes for the $V_{max}$, $EC_{50}$ plots are logarithmic (base 10).

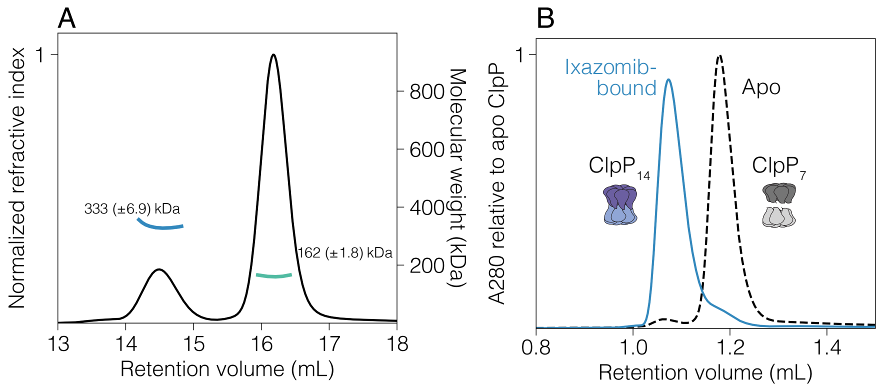


**Fig. S2.** (A) SEC-MALS demonstrates that $ClpP_{WT}$ exists as a mixture of heptamers (green trace) and tetradecamers (blue trace) in solution, with the heptameric population dominating; and (B) HPLC-based SEC shows that the addition of 10-fold molar excess of ixazomib (blue trace) results in a transition towards the tetradecameric state.

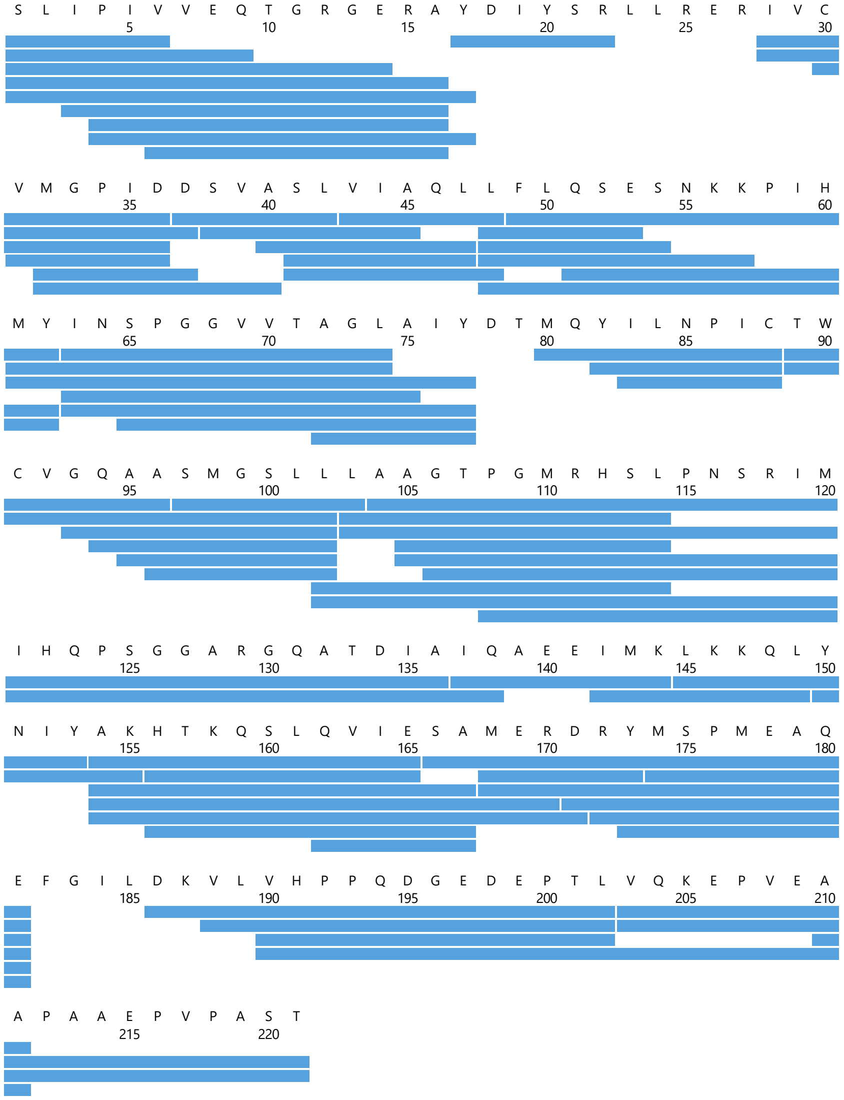

**Fig. S3.** The HDX-MS sequence coverage map of human mitochondrial ClpP, lacking its mitochondrial targeting sequence, digested online with Nepenthesin II. The Sequence coverage is 95%, and the sequence redundancy levels is 4.37.

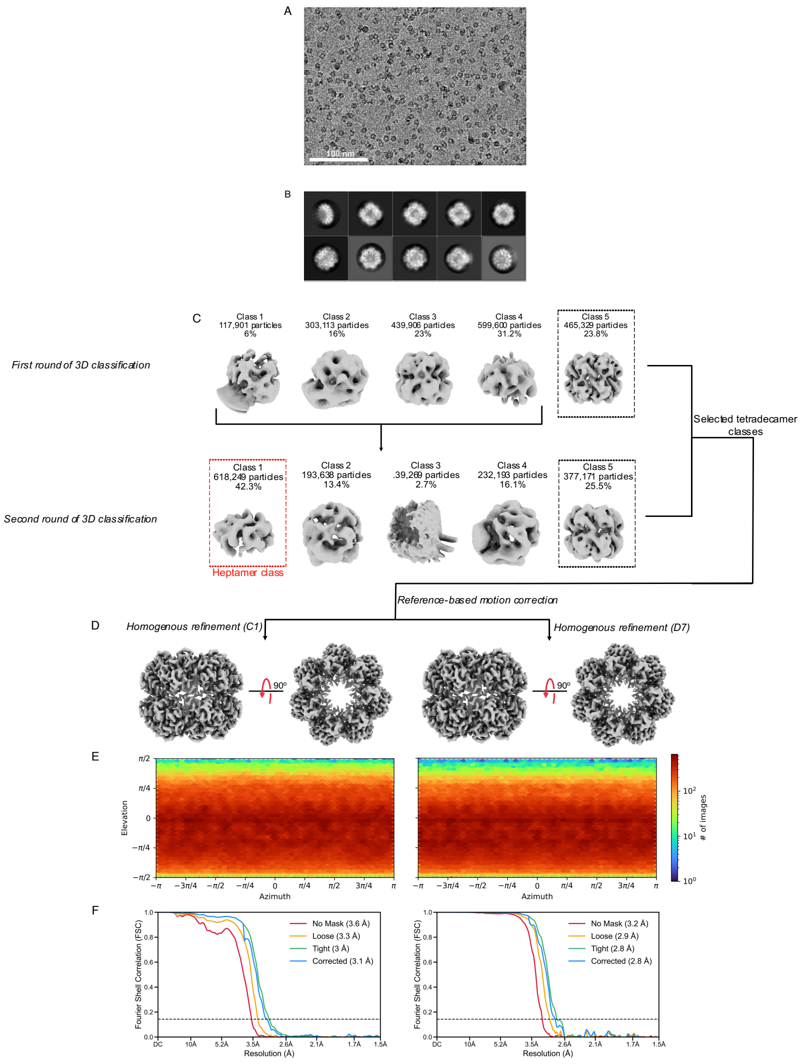


**Fig. S4. Cryo-EM data processing pipeline for apo ClpP$_{WT}$.** (A) Representative micrograph; (B) Representative 2D classes; (C) 3D classes after heterogenous refinement; (D) Final consensus maps with C1 (left) and D7 (right) symmetry imposed; (E) Viewing direction distribution plots for C1 (left) and D7 (right) refinements; and (F) Gold-standard Fourier shell correlation plots for C1 (left) and D7 (right) refinements.

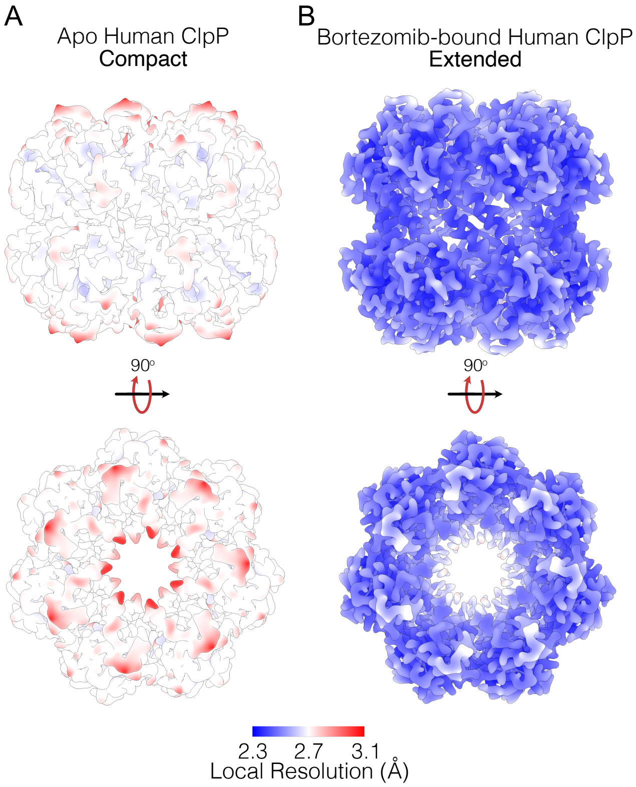


**Fig. S5.** Density maps of apo and bortezomib-bound ClpP$_{WT}$ coloured according to local resolution. Bortezomib binding results in the transition of the ClpP barrel to the activated extended state.

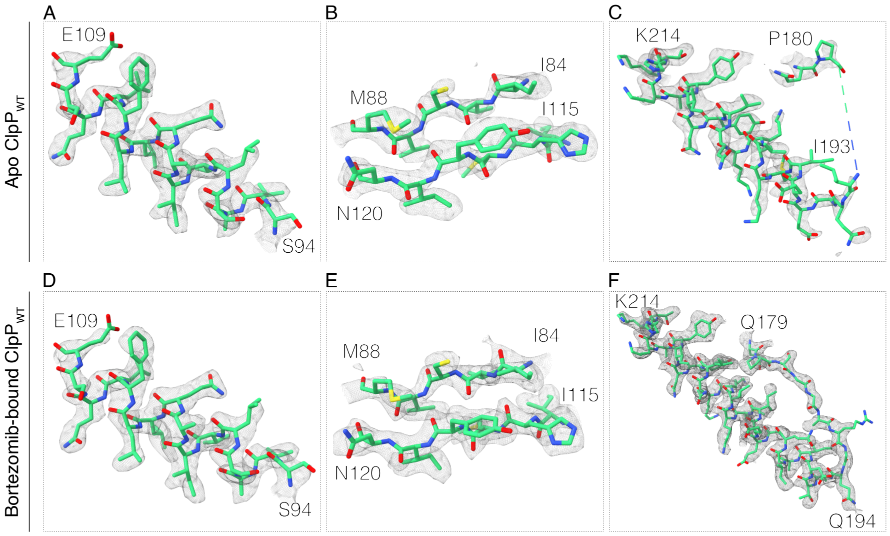


**Fig. S6. Model-in consensus map fits for selected ClpP$_{WT}$ regions.** (A) Apo ClpP$_{WT}$ helix 2, (B) β-strands 1 and 2, and (C) handle helix. (D) Bortezomib-bound ClpP$_{WT}$ helix 2 (E) β-strands 1 and 2 (F) handle domain.

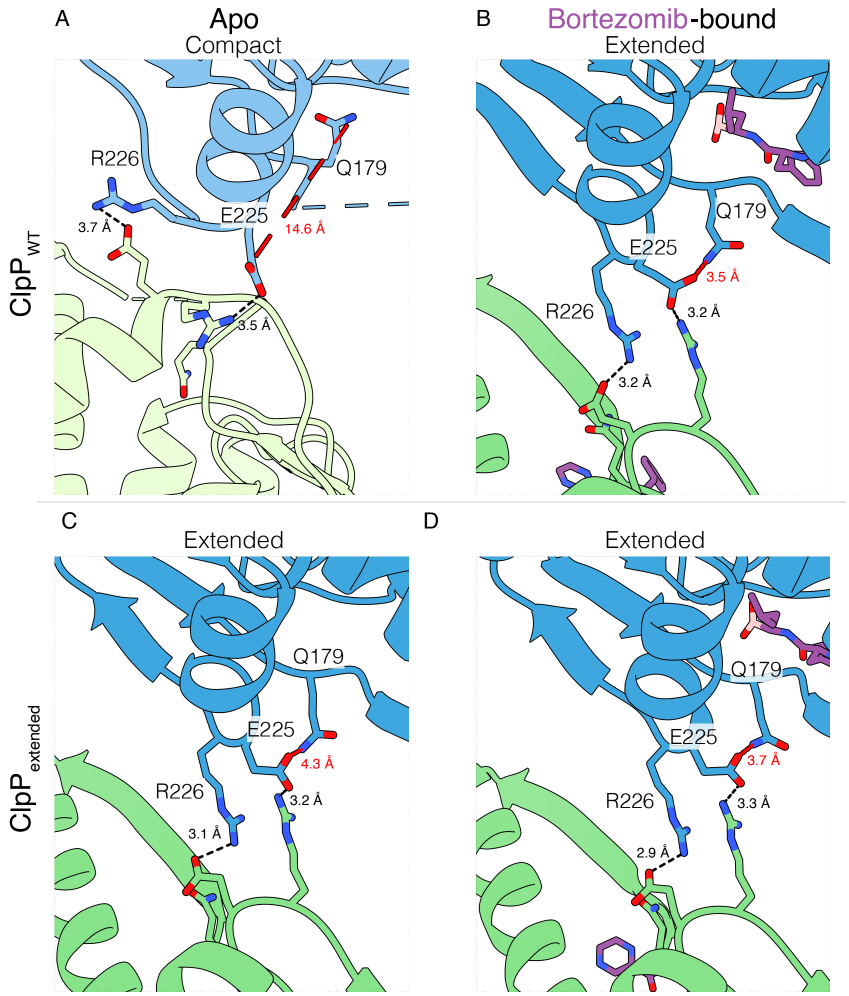


**Fig. S7. Oligomerization sensors and segment of an electrostatic network critical for ClpP function.** (A) The structure of apo $ClpP_{WT}$ is in the compact state, where the oligomerization residues (E225 and R226) are close enough to interact (3.5 and 3.7Å), however Q179, which is critical for positioning the catalytic His faces the interior of the barrel 14.6 Å away from E225; (B) Bortezomib-bound $ClpP_{WT}$; (C) Apo $ClpP_{extended}$ and (D) Bortezomib-bound $ClpP_{extended}$ are in the extended state representative by engaged oligomerization sensors and electrostatic network (distances <4 Å ).

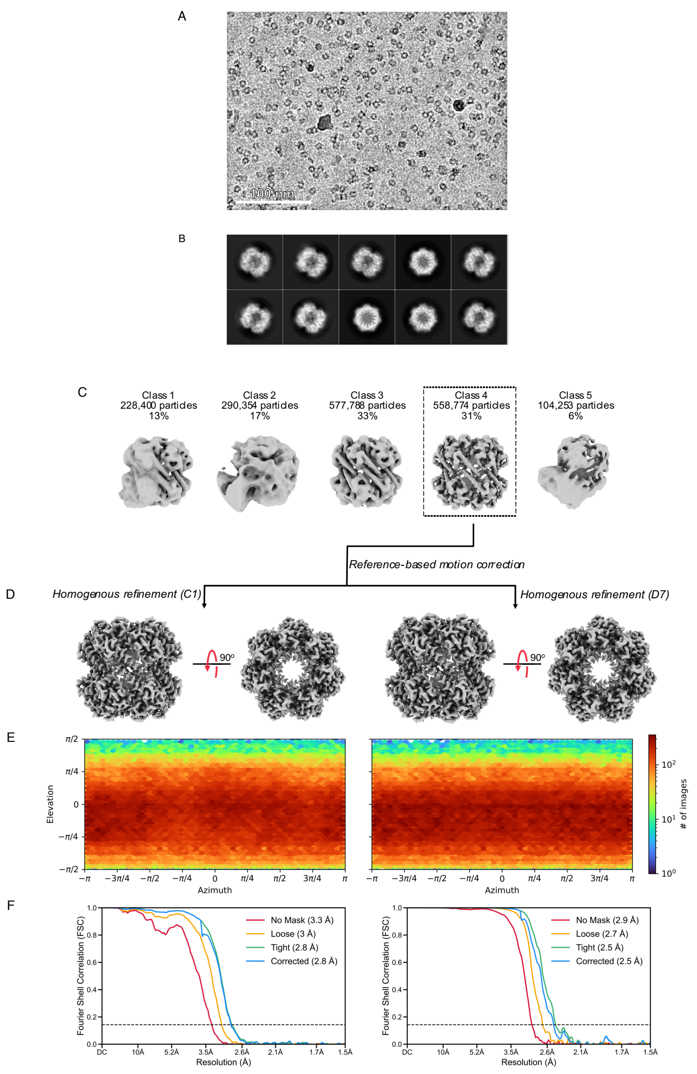


**Fig. S8. Cryo-EM data processing pipeline for bortezomib-bound $ClpP_{WT}$.** (A) Representative micrograph; (B) Representative 2D classes; (C) 3D classes after heterogenous refinement; (D) Final consensus maps with C1 (left) and D7 (right) symmetry imposed; (E) Viewing direction distribution plots for C1 (left) and D7 (right) refinements; and (F) Gold-standard Fourier shell correlation plots for C1 (left) and D7 (right) refinements.

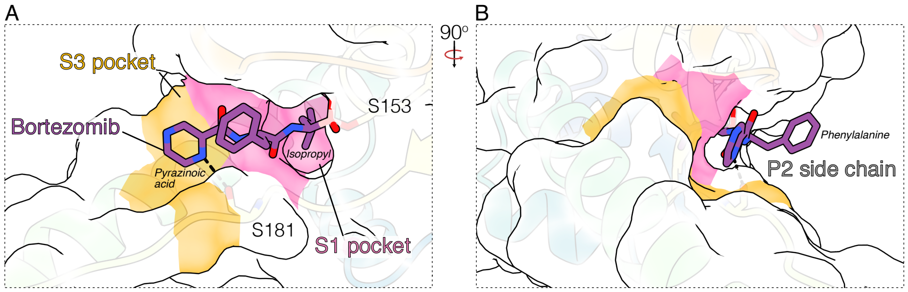


**Fig. S9. Surface representation of the active site of bortezomib-bound ClpP$_{WT}$ structure.** (A) The isopropyl moiety of bortezomib (purple) sits in the deep non-polar S1 pocket lined by the side chains of V126, M154, and P180, Y209, and M224 (pink). The pyrazinoic acid group sits shallowly in the S3 pocket formed by G182, I198, and L201 (orange) and interacts with the backbone carbonyl of S181; and (B) Human ClpP lacks an S2 pocket, therefore the central phenylalanine moiety of bortezomib is largely exposed on the protein surface.

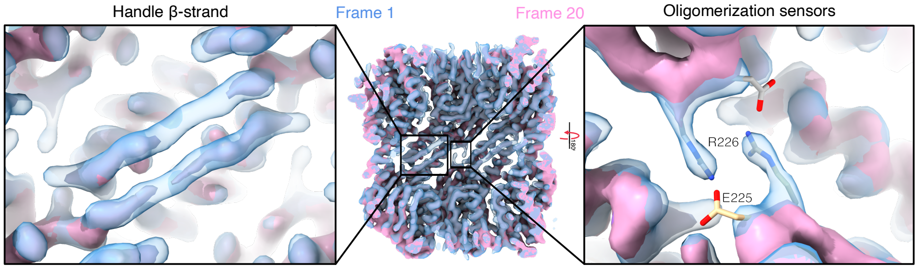


**Fig. S10.** Three-dimensional variability analysis (3DVA) analysis of bortezomib-bound $ClpP_{WT}$, showing an overlay of maps representing the first (blue) and last (pink) frames of a component that visualizes the transient formation and disappearance of density corresponding to the handle β-strand and oligomerization sensor residues.

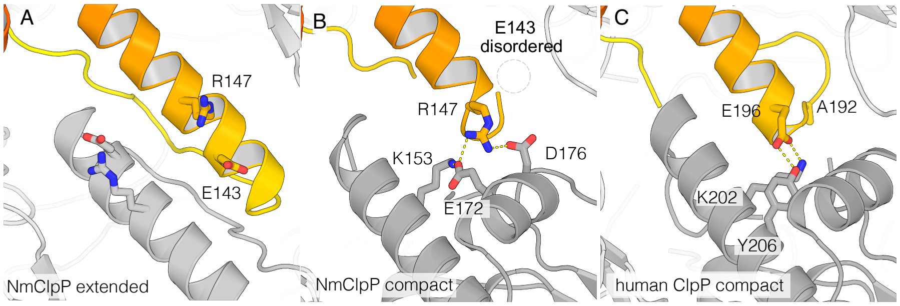


**Fig. S11.** (A) Organization of E143 and R147 in the handle domain of *Neisseria meningitis* ClpP in the extended conformation (PDB 6NAQ). These residues form an electrostatic pair that help stabilize the handle domain; (B) Organization of E143 and R147 in *Neisseria meningitidis* ClpP in the compact conformation. Here, R147 participates in a network of favorable hydrogen bonds and electrostatic interactions with the partner protomer from the opposite ring, while E143 is disordered in the lumen. (C) Organization of E196 and A192 in the compact state of wild-type human ClpP (PDB 6BBA). E196 partakes in favourable charge-charge interactions and hydrogen bonds with K202 and Y206 from the protomer in the opposite ring, helping stabilize the compact conformation (PDB 6BBA).

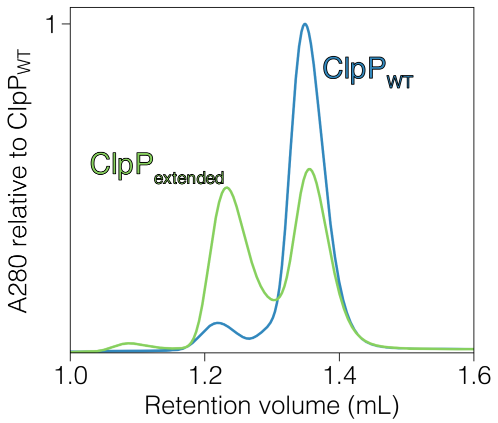


**Fig. S12.** SEC reveals that the A192E / E196R mutations in $ClpP_{extended}$ significantly shift the equilibrium towards the tetradecameric state compared to $ClpP_{WT}$. The running buffer for SEC was 50 mM Tris – pH 7.8, 200 mM KCl, 10 mM $MgCl_2$ and 1 mM DTT.

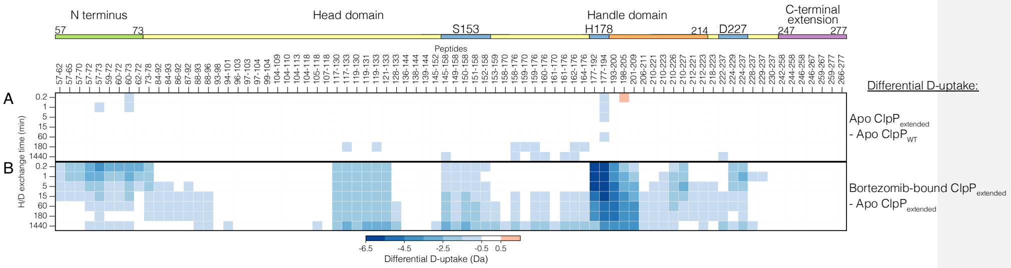


**Fig. S13.** (A) A heat map displaying the deuterium uptake of apo $ClpP_{extended}$ relative to $ClpP_{WT}$, showing rigidification on a handle domain peptide (residues 177 – 194). (B) The deuterium uptake of bortezomib-bound $ClpP_{extended}$ relative to apo $ClpP_{extended}$ follows a similar deuterium uptake reduction pattern to bortezomib-bound $ClpP_{WT}$ with the most significant protection observed on N-terminal and handle domain peptides.

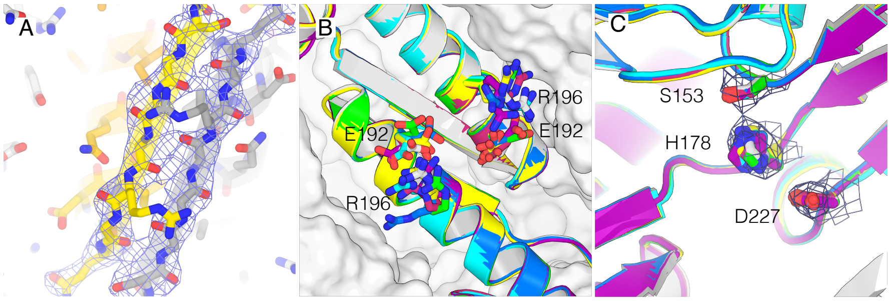


**Fig. S14.** (A) X-ray electron density contoured at 1.2 σ around the β-strand of the handle domain of apo $ClpP_{extended}$; (B) Details of packing the engineered intrachain salt bridge between E192 and R196, with each of the seven independent chain pairs superimposed and shown in a different colour. The density for the sidechains of these residues is generally weak, suggesting residual mobility, but they do approach one another close enough to form a hydrogen bond in about half of the chains; and (C) Details showing the organization of the active site. Electron density is shown for one chain, contoured at 1σ.

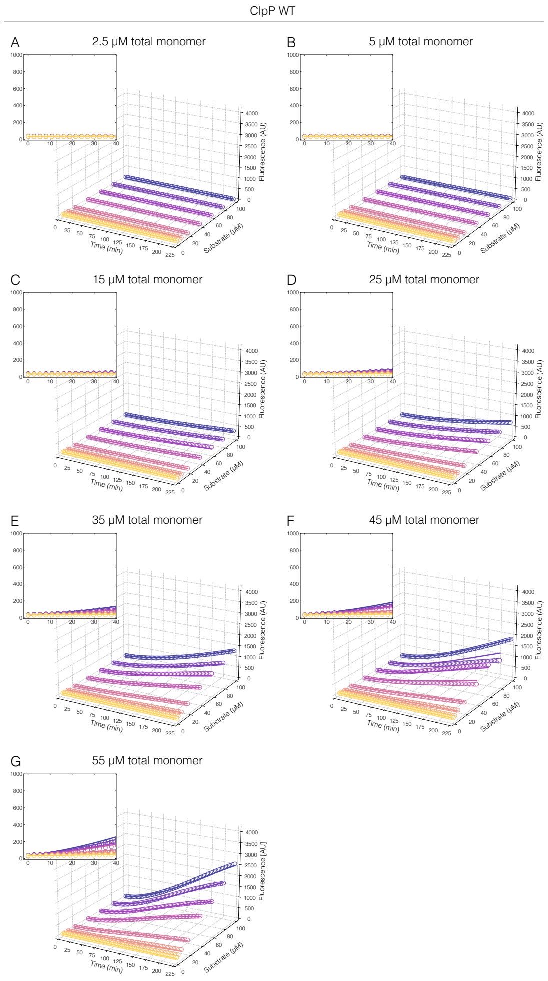


**Fig. S15.** Global fits of $ClpP_{WT}$ peptidase activity profiles. (A-G) Reaction progress curves monitoring increases in fluorescence from cleavage of a reporter peptide by $ClpP_{WT}$ are shown as coloured circles; only every third data point is displayed for clarity. Total $ClpP_{WT}$ monomer concentrations of 2.5, 5, 15, 25, 35, 45, and 55 µM were implemented in individual reactions measured with substrate concentrations of 2.5, 5, 10, 20, 40, 60, 80, and 100 µM. Insets highlight the non-linear nature of the activity curves on the timescale of 0 to 40 minutes. Global fits of the kinetic model in Figure 4C top (see Supplementary Materials and Methods for details) are displayed as coloured curves.

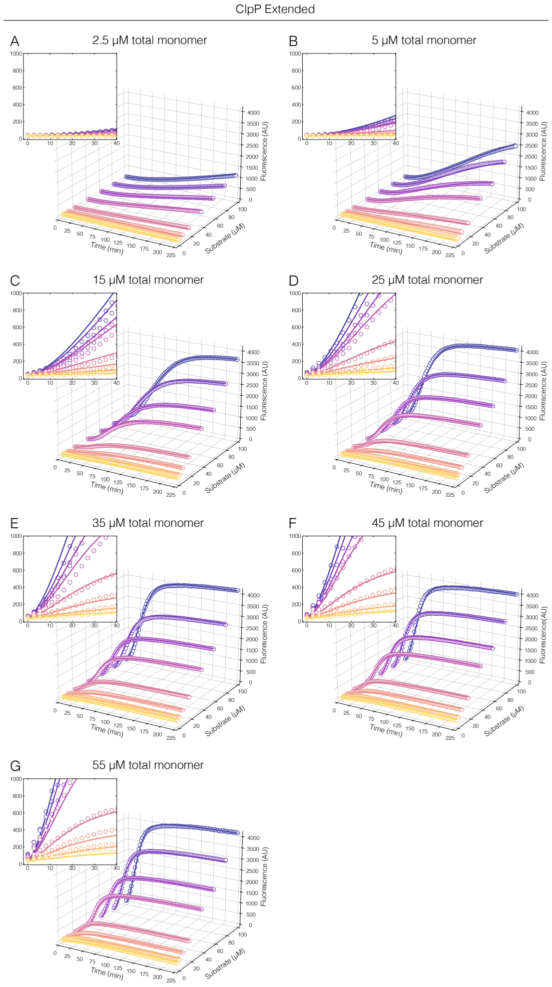


**Fig. S16.** Global fits of ClpP$_{extended}$ peptidase activity profiles. (A-G) Reaction progress curves monitoring increases in fluorescence from cleavage of a reporter peptide by ClpP$_{extended}$ are shown as coloured circles; only every third data point is displayed for clarity. Total ClpP$_{extended}$ monomer concentrations of 2.5, 5, 15, 25, 35, 45, and 55 µM were implemented in individual reactions measured with substrate concentrations of 2.5, 5, 10, 20, 40, 60, 80, and 100 µM. Insets highlight the curved nature of the reaction progress profiles on the timescale of 0 to 40 minutes. Global fits of the kinetic model in Figure 4C top (see Supplementary Materials and Methods for details) are displayed as coloured curves.

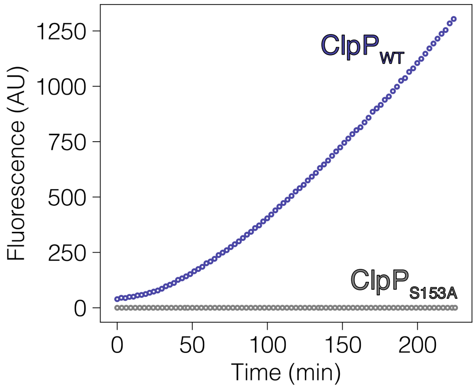


**Fig. S17.** Mutation of catalytic S153 to alanine demonstrates that all activity in the protein samples originated from human mitochondrial ClpP. Reaction progress curves recorded on ClpP$_{WT}$ (purple) and ClpP$_{S153A}$ (grey) at 35 $\mu$M with Ac-WLA-AMC at 100 $\mu$M.

### SI Appendix Movies

**Movie S1**. 3DVA performed on apo ClpP$_{WT}$ displaying the appearing and disappearing of density for the oligomerization sensors.

**Movie S2**. 3DVA performed on bortezomib-bound ClpP$_{WT}$ displaying the appearing and disappearing of density for the handle β-strand and oligomerization sensors.

**Movie S3**. 3DVA performed on bortezomib-bound ClpP$_{WT}$ displaying flexibility in the N-terminal β-hairpin.

**Table S1. HDX summary.**

| | Apo $ClpP_{WT}$ | Bortezomib-bound $ClpP_{WT}$ | Apo $ClpP_{extended}$ | Bortezomib-bound $ClpP_{extended}$ |
|---|---|---|---|---|
| HDX reaction details | Final $D_2O$ concentration (v/v): 90% $pH_{corr}$ 7.8, RT 4% DMSO | Final $D_2O$ concentration (v/v): 90% $pH_{corr}$ 7.8, RT 2 mM Bortezomib 4% DMSO | Final $D_2O$ concentration (v/v): 90% $pH_{corr}$ 7.8, RT 4% DMSO | Final $D_2O$ concentration (v/v): 90% $pH_{corr}$ 7.8, RT 2 mM Bortezomib 4% DMSO |
| HDX time course (min) | 0.16, 1, 5, 15, 60, 180, 1440 | | | |
| Undeuterated controls | 3 | | | |
| Back-exchange | 33 % (estimated from Turner, M. *et al* 2024 (27)) | | | |
| Number of peptides | 81 | | | |
| Sequence coverage | 95% | | | |
| Average peptide length/redundancy | 11.3 ± 4.1 / 4.37 | | | |
| Replicates | 3 technical replicates | | | |
| Repeatability | 0.14 Da | | | |
| Significant differences | 0.5 Da | | | |

**Commented [MG2]:** Cite, not sure how to cite bioRxiv

**Table S2: Data collection and refinement statistics for cryo-EM**

| | Apo $ClpP_{WT}$ | Bortezomib-bound $ClpP_{WT}$ |
|---|---|---|
| PDB ID | **9DKV** | **9DKW** |
| EMDB ID | **EMD-46970** | **EMD-46971** |
| **Data collection and processing** | | |
| Microscope and camera | FEI Titan Krios with Gatan K3 Camera | FEI Titan Krios with Gatan K3 Camera |
| Magnification (x) | 130,000 | 130,000 |
| Voltage (kV) | 300 | 300 |
| Data acquisition software | Serial EM | Serial EM |
| Electron dose (e-/A2) | 60 | 60 |
| Defocus range (um) | -1.00 to 2.50 | -1.00 to 2.50 |
| Pixel size (A) | 0.675 | 0.675 |
| Number of micrographs | 6615 | 6210 |
| Processing software | cryoSPARC v4 | cryoSPARC v4 |
| Symmetry imposed | D7 | D7 |
| Initial particle images (no.) | 1,997,332 | 1,995,006 |
| Final particle images (no.) | 768,999 | 408,161 |
| Map resolution (at FSC=0.143) (A) | 2.81 | 2.49 |
| **Refinement** | | |
| Intial model used | PDB ID : 8I7X | PDB ID : 8I7X |
| Model resolution - Masked(FSC=0/0.143/0.5) | 2.7/2.8/2.9 | 2.4/2.4/2.6 |
| Map sharpening factor (A2) | 176.4 | 130.9 |
| **Model Composition** | | |
| Non-hydrogen atoms | 18368 | 20314 |
| Protein residues | 2310 | 2548 |
| Ligands | 0 | BO2:14 |
| **B-factors (A2) (min/max/mean)** | | |
| Protein | 0.84/86.44/28.13 | 5.34/83.41/26.53 |

| | | |
|---|---|---|
| Ligand | NA | 11.79/43.41/27.42 |
| **r.m.s. deviations** | | |
| Bond lengths (A) | 0.004 | 0.003 |
| Bond angles (deg) | 0.588 | 0.712 |
| **Model Validation** | | |
| Molprobity score | 1.11 | 1.14 |
| Clash score | 3.17 | 3.45 |
| Poor rotamers (%) | 0.5 | 1.01 |
| **Ramachandran plot** | | |
| Favoured (%) | 97.98 | 99.2 |
| Allowed (%) | 2.02 | 0.8 |
| Dissalowed (%) | 0 | 0 |

**Table S3. Data collection and refinement statistics.**

| | Apo ClpP$_{Extended}$ | Bortezomib-bound ClpP$_{Extended}$ |
|---|---|---|
| **PDB ID** | 9DQK | 9DQL |
| **Wavelength** | 0.9536 | 0.9536 |
| **Resolution range** | 49.07 - 2.75 (2.848 - 2.75) | 44.15 - 3.2 (3.314 - 3.2) |
| **Space group** | P 1 21 1 | P 1 21 1 |
| **Unit cell** | 121.32 94.99 134.36 90 96.111 90 | 122.19 93.88 133.72 90 97.874 90 |
| **Total reflections** | 545758 (55639) | 338500 (31807) |
| **Unique reflections** | 79133 (7831) | 49648 (4912) |
| **Multiplicity** | 6.9 (7.1) | 6.8 (6.5) |
| **Completeness (%)** | 99.87 (99.95) | 99.68 (99.92) |
| **Mean I/sigma(I)** | 13.00 (1.18) | 11.19 (1.21) |
| **Wilson B-factor** | 82.26 | 123.28 |
| **R-merge** | 0.09889 (1.726) | 0.1256 (1.851) |
| **R-meas** | 0.107 (1.863) | 0.136 (2.016) |
| **R-pim** | 0.04055 (0.6954) | 0.05171 (0.7894) |
| **CC1/2** | 0.999 (0.465) | 0.998 (0.402) |
| **CC*** | 1 (0.797) | 1 (0.757) |
| **Reflections used in refinement** | 79092 (7831) | 49620 (4911) |
| **Reflections used for R-free** | 1730 (171) | 1696 (168) |
| **R-work** | 0.2063 (0.3744) | 0.1860 (0.3586) |
| **R-free** | 0.2504 (0.4040) | 0.2365 (0.3976) |

| | | |
|---|---|---|
| **CC(work)** | 0.966 (0.610) | 0.976 (0.584) |
| **CC(free)** | 0.942 (0.506) | 0.945 (0.519) |
| **Number of non-hydrogen atoms** | 19762 | 20087 |
| **macromolecules** | 19686 | 19686 |
| **ligands** | 14 | 401 |
| **solvent** | 62 | 0 |
| **Protein residues** | 2538 | 2535 |
| **RMS(bonds)** | 0.002 | 0.003 |
| **RMS(angles)** | 0.43 | 0.49 |
| **Ramachandran favored (%)** | 96.29 | 96.05 |
| **Ramachandran allowed (%)** | 3.63 | 3.79 |
| **Ramachandran outliers (%)** | 0.08 | 0.16 |
| **Rotamer outliers (%)** | 1.01 | 4.7 |
| **Clashscore** | 4.2 | 7.69 |
| **Average B-factor** | 94.51 | 132.5 |
| **macromolecules** | 94.55 | 132.55 |
| **ligands** | 92.86 | 130.4 |
| **solvent** | 81.66 | |
| **Number of TLS groups** | 83 | 90 |

Statistics for the highest-resolution shell are shown in parentheses.
